## Supplementary material for "Eco-evolutionary dynamics of nested Darwinian populations and the emergence of community-level heredity": Python code, figures and simulations: escaffolding.py

```
""" escaffolding.py -- functions to perform ecological saffolding.
This file is part of the ecological scaffolding package.
Copyright 2018 Guilhem Doulcier, Licence GNU GPL3+
"""

import multiprocessing
import pickle
import numpy as np
import logging

logger = logging.getLogger('estaudel')
logger.setLevel(logging.DEBUG)
logger.propagate = False
ch = logging.StreamHandler()
ch.setLevel(logging.DEBUG)
logger.addHandler(ch)


def load(file):
    """Create a new Output object and load its content from a pickle file"""
    out = Output(0, 0, 0, 0)
    err = out.load(file)
    if not err:
        return out
    else:
        raise IOError


class Output:
    """ Store the output from the simulation.

    This object handle the saving and loading of the simulation data.
    The method append is called each generation and fill the attributes.

    Attributes:
        phenotype[i,j,n,d]: Trait j value of type i in collective d at generation n.
        fitness[n,d]: fitness of collective d at genration n
        state[i,n,d]: Number of individuals of type i in collective d at generation n.
        parents[n][d]: Identity of the parent of collective d in generation n+1.
        parameters : dictionnary of parameters
        data: dictionnary used to store additional information deduced from the other attributes.

    """

    def __init__(self, N: int, D: int, Ntypes: int, Psize: int):
        """
        Args:
        - Ntypes: number of different phenotypes in the population.
        - Psize: size of a phenotype .
        - N: Number of generations.
        - D: Number of collectives.
        """

        # State of all the collectives:
        # All the array are assigned empty at the full size for performance reason.
        self.phenotype = np.empty((Ntypes, Psize, N, D))
        self.fitness = np.empty((N, D))
        self.state = np.empty((Ntypes, N, D))
        self.parents = []

        # Current generation.
        self.current_gen = 0

        self.parameters = {}
        self.data = {}

    def append(self, gen: int, icol: int, out: object):
        """Called at each collective generation `gen` for collective `icol` with
        `out` the output of the growth function.
        """
        self.phenotype[:, :, gen, icol] = out['phenotype']
        self.fitness[gen, icol] = out['fitness']
        self.state[:, gen, icol] = out['state']
        self.current_gen = gen

    def __repr__(self):
        return """Ecological scaffolding data {} generations""".format(self.current_gen)

    def save(self, filename):
        """Save the content of the object in a pickle file"""
        with open(filename, 'wb') as file:
            pickle.dump({'phenotype': self.phenotype,
                         'fitness': self.fitness,
                         'state': self.state,
                         'current_gen': self.current_gen,
                         'parents': self.parents,
                         'parameters': self.parameters,
                         'data': self.data
                         }, file)

    def load(self, filename):
        """Load the content of the object from a pickle file"""
        with open(filename, 'rb') as file:
            try:
                data = pickle.load(file)
            except OSError as ex:
                logger.warning('OS Error loading {} {}'.format(filename, ex))
                return 1
            except EOFError as ex:
                logger.warning('File Error loading {} {}'.format(filename, ex))
                return 1
            else:
                self.phenotype = data['phenotype']
                self.fitness = data['phenotype']
                self.state = data['state']
                self.parents = data['parents']
                self.current_gen = data['current_gen']
                self.parameters = data['parameters']
                self.data = data['data'] if 'data' in data else {}
                return 0


def collective_generations(N: int, pop, output,
                           collective_fitness_func,
                           collective_birth_death_process_func,
                           dilution_and_growth_func, filename=None, save_frequency=100, pool=None):
    '''Args:
        N: number of generations
        pop: list of <phenotype, state>
        output: object, its append method is called each generation.
        collective_fitness_func: collective fitness of a <phenotype, state>
        collective_birth_death_process_func: return the list of parents from a list of fitness
        dilution_and_growth_func: transform pop.
    '''
    if pool is None:
        pool = multiprocessing.Pool(1)
    for n in range(output.current_gen, output.current_gen+N):
        logger.info('Generation {}'.format(n))
        grown = pool.map(dilution_and_growth_func, pop, 1)
        for d, out in enumerate(grown):
            out['fitness'] = collective_fitness_func(out['phenotype'], out['state'])
            output.append(n, d, out)
        parents = collective_birth_death_process_func([col['fitness'] for col in grown])
        if not len(parents):
            logger.warning('No surviving collective')
            break
        output.parents.append(parents)
        pop = [(grown[i]['phenotype'], grown[i]['state']) for i in parents]
        if filename is not None and n % save_frequency == 0:
            fname = '{}_ongoing.pkle'.format(filename)
            logger.debug('Saving to {}'.format(fname))
            output.save(fname)
    return output, pop


def collective_serial_transfer(fitness, **_):
    """Decide wich collective should be reproduced and which should be
    discarded. All collective have exactly one child here.

    Return:
        a list containing the indice of the parent of each new collective
    """
    return list(range(len(fitness)))


def collective_birth_death_neutral(fitness, percentile=None):
    """Decide wich collective should be reproduced and which should be
    discarded.

    Here we discard a percentile of collectives chosen unifromly.
    Args:
        fitness (iterable): fitness of each collective.
        percentile (float): percentile of collectives to discard.
    Return:
        a list containing the indice of the parent of each new collective
    """
    if percentile is None:
        return collective_serial_transfer(fitness)

    D = len(fitness)
    nborn = int(D * percentile/100.0)
    nsurv = D-nborn
    surviving = list(np.random.choice(np.arange(len(fitness)), size=nsurv, replace=False))
    logger.info('Percentile is {}, {} survivors '.format(percentile, len(surviving)))
    newborns = list(np.random.choice(surviving, size=nborn))
    return surviving + newborns


def collective_birth_death_process(fitness, percentile=None):
    """Decide wich collective should be reproduced and which should be
    discarded.

    Args:
       fitness (iter): fitness of each parent collective.
    Return:
        a list containing the indice of the parent of each new collective
    """
    fitness = np.array(fitness)

    # If all fitness are equal, neutral
    if all(fitness == np.max(fitness)):
        logger.warning('All collective have the same fitness...')
        return collective_birth_death_neutral(fitness, percentile)

    if percentile is None:
        surviving = [d for d, f in enumerate(fitness) if np.random.random() < f]
    else:
        threshold = np.percentile(fitness, percentile)
        surviving = [d for d, f in enumerate(fitness) if f > threshold]
        logger.info('Percentile is {}, fitness threshold is {} '.format(
            percentile, threshold))
    ndeath = len(fitness)-len(surviving)

    logger.info('{} Collective Death ~ Mean fitness {}'.format(ndeath, np.mean(fitness)))

    # if no survivor, return an empty list
    if not surviving:
        return surviving

    # Newborns are taken uniformely from the surviving individuals to
    # keep pop size constant.
    newborns = list(np.random.choice(surviving, size=ndeath))

    return surviving+newborns


def collective_birth_death_process_soft(fitness, percentile):
    """Decide wich collective should be reproduced and which should be
    discarded.

    Only `percentiles` of collectives survive at each generation,
    following a weighted-by-fitness drawing without replacement.
    Collective population size is kept constant by drawing uniformly with replacement the
    parents of the next collective generation among the surviving collectives.

    Args:
       fitness (iter): fitness of each parent collective. Weight of the drawing.
       percentiles (float): proportion (in %) of collectives to discard.
    Return:
        a list containing the indice of the parent of each new collective.
    """
    fitness = np.array(fitness)
    D = len(fitness)
    nborn = int(D * percentile/100.0)
    nsurv = D-nborn

    # If all fitness are equal, default to neutral
    if all(fitness == np.max(fitness)):
        return collective_birth_death_neutral(fitness, percentile)

    surviving = list(np.random.choice(np.arange(len(fitness)),
                                      size=nsurv,
                                      replace=False,
                                      p=fitness/fitness.sum()))

    logger.info('{} Collective Death ~ Mean fitness {}'.format(nborn, np.mean(fitness)))

    # Newborns are taken uniformely from the surviving individuals to
    # keep pop size constant.
    newborns = list(np.random.choice(surviving, size=nborn))
    return surviving+newborns


def dilution_and_growth(parent, bottleneck_size, growth_function):
    """Perform dilution then growth of a collective

    Args:
        parent (state,phenotypes).
        bottleneck size (int): number of cells.
        growth_function (function): perform a growth.
    """
    state = np.random.multinomial(bottleneck_size, parent[1]/parent[1].sum())
    return growth_function(state=state, phenotypes=parent[0])
```
