## Supplementary material for "Eco-evolutionary dynamics of nested Darwinian populations and the emergence of community-level heredity": Python code, figures and simulations: deterministic.py

```
""" heredity/deterministic.py -- Evolution of heredity thought experiment, deterministic model.
This file is part of the ecological scaffolding package/ heredity model subpackage.
Copyright 2018 Guilhem Doulcier, Licence GNU GPL3+
--
"""
from functools import partial
import numpy as np
import scipy.integrate

def lotka_volterra(x, _=None, r=None, a=None):
    '''Temporal derivative of the Lotka-Volterra system.
    Args:
        x (array of N floats): number of individual of each type.
        _ (None): time, not used but required by scipy.integrate.
        r (array of N floats): maximum growth rate of each type.
        a (array of NxN floats): inter-type interaction coefficient.

    Return dx/dt.
    '''
    return (r*x)*(1-a@x)

def lotka_volterra_Nf(x,_=None,r=None,a=None):
    '''Temporal derivative of the Lotka-Volterra system in Nf coordinates.
    Args:
        x (array of 2 floats): x[0] = N (number of individuals, x[1] = f (frequency of type 1 individuals).
        _ (None): time, not used but required by scipy.integrate.
        r (array of N floats): maximum growth rate of each type.
        a (array of NxN floats): inter-type interaction coefficient.

    return (dN/dt, df/dt)'''
    return np.array([
     (r[0]*(-a[0,0]*x[0]*x[1] + a[0,1]*(x[1] - 1)*x[0] + 1)*x[1] - r[1]*(x[1] - 1)*(-a[1,0]*x[0]*x[1] + a[1,1]*(x[1] - 1)*x[0] + 1))*x[0],
     (r[0]*(a[0,0]*x[0]*x[1] - a[0,1]*(x[1] - 1)*x[0] - 1) - r[1]*(a[1,0]*x[0]*x[1] - a[1,1]*(x[1] - 1)*x[0] - 1))*(x[1] - 1)*x[1]
    ])

def iterate_ecology(r, A, B, T, tstep=500, generations=8, x0=0.1):
    """Returns the ecological trajectory across several generations.

    r (array of 2 floats): maximum growth rate of each type.
    A (array of 2x2 floats): inter-type interaction coefficient.
    B (float): Bottleneck size.
    T (float): Duration of growth rate
    tstep (int): number of integration steps per generation.
    generations (int): number of collective generations
    x0 (float): initial proportion of types.

    Returns a list of `generations` succesive trajectories and a list
    of `generations`+1 final proportions
    """
    flow = partial(lotka_volterra, r=r, a=A)
    time = np.linspace(0, T, tstep)
    plist = [x0]
    traj = []

    x = x0
    for _ in range(generations):
        xy = scipy.integrate.odeint(flow,
                                    y0=np.array([B*x, B*(1-x)]),
                                    t=time)
        x = float(xy[-1, 0]/xy[-1, :].sum())
        traj.append(xy)
        plist.append(x)

    return traj, plist

def convert_phenotypes_to_lv(phenotypes, K):
    """Convert the parameters of the stochastic model to the parameters of
    the deterministic model.

    Return the growth array r and interaction matrix A."""
    try:
        phenotypes.shape[0]
    except:
        phenotypes = np.array(phenotypes)

    if phenotypes.shape[0] == 2:
        pos_0 = list(phenotypes[:, 0]).index(0)
        pos_1 = list(phenotypes[:, 0]).index(1)
        r = np.array([phenotypes[pos_0, 1], phenotypes[pos_1, 1]])
        A = np.array([[phenotypes[pos_0, 2]/K, phenotypes[pos_1, 3]/K],
                      [phenotypes[pos_0, 3]/K, phenotypes[pos_1, 2]/K]])
    else:
        r = np.array(phenotypes[:, 1])
        A = np.empty(shape=(phenotypes.shape[0],phenotypes.shape[0]))
        for i, focal in enumerate(phenotypes):
            A[i,i] = focal[2]
            for j, other in enumerate(phenotypes):
                A[i,j] = other[2+int(focal[0] != other[0])]
        A /= K
    return r,A

def internal_equilibrium(A):
    det = np.linalg.det(A)
    stable = A[0, 0] > A[1, 0] and A[1, 1] > A[0, 1]
    return [(A[1, 1]-A[0, 1])/det, (A[0, 0]-A[1, 0])/det], stable

def stable_equilibria(A):
    """Output the stable equilibrium point provided with the interaction
    matrix."""
    if A[0, 0] > A[1, 0] and A[1, 1] > A[0, 1]:
        # Coexistence
        return internal_equilibrium(A)
    elif  A[0, 0] > A[1, 0]:
        # 0 exclude 1
        return [1/A[0, 0], 0]
    elif  A[0, 0] > A[1, 0]:
        # 1 exclude 0
        return [0, 1/A[1, 1]]
    # Bistable.
    return [np.nan,np.nan]

def list_equilibria(A):
    """List equilibria associated with the interaction matrix"""
    ie, ie_stability = internal_equilibrium(A)
    equilibria = [[0, 0], [0, 1/A[1, 1]], [1/A[0, 0], 0], ie]
    stability = [False,
                 A[0, 1] > A[1, 1],
                 A[1, 0] > A[0, 0],
                 ie_stability]
    return list(zip(equilibria, stability))

def pstar(A):
    '''Return the proportion of type 1 cells at the stable
    attractor'''
    if A[0, 0] > A[1, 0] and A[1, 1] > A[0, 1]:
        # Coexistence
        return (A[1, 1] - A[0, 1])/(A.trace()-A[1, 0]-A[0, 1])
    elif  A[0, 0] > A[1, 0]:
        # 0 exclude 1
        return 0
    elif  A[1, 1] > A[0, 1]:
        # 1 exclude 0
        return 1
    # Bistable.
    return np.nan

def tstar(r, A, B, precise=False, steps=100, eps=1e-6):
    '''Return the value of T where an internal fixed point appear.

    Args:
        r (np.array, size 2): growth rate
        A (np.array, size 2x2): interaction matrix
        B (int): bottleneck size
        precise (bool): see below
        steps (int): max number of steps for the numerical approx.
        eps (float): precision for the numerical approx.

    If precise is False, will only output a rough analytical
    approximation. Otherwise, will try to numerically compute it.

    Returns np.nan if there is no internal fiexd point.
    '''
    stable_eq = pstar(A)

    ifast, islow = (1, 0) if r[1] > r[0] else (0, 1)


    # If the fast growing type is also the one excluding the other,
    # there is no critical time.
    if ((ifast == 0 and np.isclose(stable_eq, 1))
        or (ifast == 1 and np.isclose(stable_eq, 0))):
        return np.nan, np.nan

    guesstimate = (((1/r[ifast])*(A[islow, ifast]/A[ifast, ifast]) - (1/r[islow]))
                   / (1 - (A[islow, ifast]/A[ifast, ifast]))
                   * np.log(B*A[ifast, ifast]/(B*A[ifast, ifast]+1)))
    if not precise:
        return guesstimate, guesstimate

    low = 0+eps
    high = 1-eps
    gfunc = get_gfunc(r, A, B)
    negative_if_fp = lambda t: (gfunc(low, t)-low) * (gfunc(high, t)-high)

    return scipy.optimize.fsolve(negative_if_fp, guesstimate), guesstimate

def get_gfunc(r, A, B, T=None, tstep=1000):
    """ Return the G function (final proportion as a function of initial proportions)
    for a given set of parameters

    Args:
        r (np.array, size 2): growth rate
        A (np.array, size 2x2): interaction matrix
        B (int): bottleneck size
        T (float): duration of the growth phase
        tstep (int): integration setps.

    Return: function (float in [0,1]) -> (float in [0,1])
    """
    func = partial(lotka_volterra, r=r, a=A)

    if A is None:
        def gfunc(x,T=T):
            return 1/( 1 + ((1/x) - 1 ) * np.exp(-(T* (r[0]-r[1]))))
    else:
        def gfunc(x, T=T):
            if not T:
                return float(x)
            x = float(x)
            T = float(T)    

            xy = scipy.integrate.odeint(func,
                                        y0=np.array([B*x, B*(1-x)]),
                                        t=np.linspace(0, T, tstep))
            return float(xy[-1, 0]/xy[-1, :].sum())
    return gfunc

def continuation_on_T(gfunc, start, t0=1, tf=1e-3, tstep=100):
    """Use natural parameter continuation to draw the bifurcation diagram
    of the fixed point of G, x*, as a function of T.

    Args:
        gfunc (func: x,T -> x* [0,1]): proportion at the end of the growth
              phase as a function of the proportion at the begining of
              the growth phase (x) and the duration of the growth phase T.
       start (float): initial value of x* for T=t0
       t0 (float): initial value of T along the continuation
       tf (float): final value of T along the continuation
       tstep (int): number of values of T to compute between t0 and tf.

  Returns: list of T, list of x*
    """
    equilibria = []
    t_list = np.linspace(t0, tf, tstep)

    for T in t_list:
        fixed_point_function = lambda x, t=T: gfunc(x,t)-x
        val,_,conv,_ = scipy.optimize.fsolve(fixed_point_function,
                                             full_output=True,
                                             x0=(equilibria[-1]
                                                 if equilibria
                                                 else start))
        equilibria.append(float(val) if conv else np.nan)
    return t_list, equilibria

def continuation_left_right(func, p0, pmin, pmax, x0, steps=100):
    """Natural parameter continuation of func(x,p)=0
    on the right up to pmax and on the left up to pmin. """

    val, _, conv, _ = scipy.optimize.fsolve(lambda x,p=p0:func(x,p0), x0,
                                                full_output=True)

    if conv:
        xstart = val[0]
    else:
        raise ValueError('Starting point not found')

    # Continuation on the left
    eq_left = []
    eq_right = []
    pspace_left = np.linspace(p0, pmin, steps)
    pspace_right = np.linspace(p0, pmax, steps)
    for p in pspace_left:
        val, _, conv, _ = scipy.optimize.fsolve(lambda x,p=p:func(x,p),
                                    eq_left[-1] if eq_left else xstart,
                                    full_output=True)
        eq_left.append(val[0] if conv else np.nan)
    for p in pspace_right:
        val, _, conv, _ = scipy.optimize.fsolve(lambda x,p=p:func(x,p),
                                                eq_right[-1] if eq_right else xstart,
                                                full_output=True)
        eq_right.append(val[0] if conv else np.nan)

    pspace = np.concatenate([pspace_left[::-1], pspace_right])
    eq = eq_left[::-1]+eq_right
    return pspace, eq

def stability_of_01(gfunc, t0=1e-6, tf=1, eps=1e-6):
    """Quickly see if 0 and 1 are stable fixed point of G.

    Useful to draw the bifurcation diagram since 0 and 1 are always fixed
    points so there is no need to do a full continuation on those points.

    Args:
        gfunc (func: x,T -> x* [0,1]): proportion at the end of the growth
              phase as a function of the proportion at the begining of
              the growth phase (x) and the duration of the growth phase T.
       start (float): initial value of x* for T=t0
       t0 (float): initial value of T along the continuation
       tf (float): final value of T along the continuation
       eps (float): numerical precision
    """
    low = 0+eps
    high = 1-eps
    stability_0 = lambda t: gfunc(low,t)-low
    stability_1 = lambda t: gfunc(high,t)-high

    stable0 = stability_0(t0) < 0
    if stability_0(t0)*stability_0(tf)<0:
        tcrit =scipy.optimize.brentq(stability_0, t0, tf)
        seg0 = [[stable0,[t0, tcrit]],[not stable0,[tcrit,tf]]]
    else:
        seg0 = [[stable0,[t0,tf]]]

    stable1 = stability_1(t0) > 0
    if stability_1(t0)*stability_1(tf)<0:
        tcrit= scipy.optimize.brentq(stability_1, t0, tf)
        seg1 = [[stable1,[t0, tcrit]],[not stable1,[tcrit,tf]]]
    else:
        seg1 = [[stable1,[t0,tf]]]


    return {0:seg0, 1:seg1}

def get_bifurcation_diagram(r, A, B, tmax=1):
    """Get bifurcation diagram for T=0...tmax."""

    # Get the g function (µ,T) -> µ_final
    g = get_gfunc(r, A, B)

    # Get the equilibria of the system (T=\infty)
    ie,ies = internal_equilibrium(A)
    stability = stability_of_01(g, t0=tmax, tf=1e-9)
    ps = pstar(A)

    ts = None
    ts_approx = None
    cont_info = None
    stability_of_continuation = None
    iep = ie[0]/np.sum(ie)

    if 0 < iep < 1:
        # If the internal equilirium is stable, look for the
        # bifurcation time t* and continuate this branch from
        # t=1 to t=tstar.
        ts, ts_approx = tstar(r, A, B, precise=True)
        x0 = scipy.optimize.brentq(lambda x: g(x,1)-x, 1e-5, 1-1e-5)
        t_cont, p_cont = continuation_on_T(g, t0=tmax, tf=ts, start=x0)
        cont_info = {'t0':1, 'tmax':1, 'tmin':ts, 'p0':x0}
        stability_of_continuation = g(x0-1e-5,1)-(x0-1e-5)>0

    elif len(stability[0])==2 and len(stability[1])==2:
        # If the stability of 0 and 1 change twice on [0,1], there is
        # an intermediate branch to continuate...
        tmin = min(stability[1][0][1][1],stability[0][0][1][1])
        tmax = max(stability[1][0][1][1],stability[0][0][1][1])
        t0 = (tmin + tmax)/2
        x0 = scipy.optimize.brentq(lambda x: g(x,t0)-x, 1e-5, 1-1e-5)
        cont_info = {'t0':t0,'tmax':tmax,'tmin':tstar,'p0':x0}
        t_cont,p_cont = continuation_left_right(lambda x,t: g(x,t)-x,
                                                pmin=tmin,
                                                pmax=tmax,
                                                p0=t0,
                                                x0=x0,
                                                steps=1000)
        stability_of_continuation = g(x0-1e-5,t0)-(x0-1e-5)>0
    else:
        # Otherwise there is no continuation to perform.
        t_cont = []
        p_cont = []

    return {'pstar':ps, 'tstar':tstar,
            'tstar_approx':ts_approx,
            'g':g, 'stability_of_01':stability,
            'cont_info':cont_info,
            'continuation':[t_cont,p_cont],
            'stability_of_continuation':stability_of_continuation}
```
