## Supplementary material for "Eco-evolutionary dynamics of nested Darwinian populations and the emergence of community-level heredity": Python code, figures and simulations: display.py

```
"""heredity/display.py -- Plotting function for simulations.

This file is part of the ecological scaffolding package/ heredity model subpackage.
Copyright 2019 Guilhem Doulcier, Licence GNU GPL3+
"""
import numpy as np
import matplotlib.pyplot as plt
from mpl_toolkits.axes_grid1 import make_axes_locatable
from matplotlib.colors import ListedColormap

def plot_density(out, ax,
                 colorbar=False, cax=None, colorbar_ticks =  [1,10,100,1000],
                 ax_zoom=None, zoom_first=None,
                 fontdict=dict(size='x-large'), title=None, xcenter = 0.41):
    """ Plot the colour density of the dataset data
    out : Output object.
    ax : matplotlib axis to plot on
    colorbar: if not None, plot a colorbar in the cax axis.
    cax: colorbar axis object
    zoom_first: if not None, expand the first 'zoom_first' generations on the ax_zoom axes
    ax_zoom: axis object for the blowout of the first generations
    """

    # Preping the data
    cp_density = out.data['cp_density'] * out.parameters['D']

    # Colormap
    zbin_cp = [-100, 1]+list(np.logspace(1, np.log10(out.parameters['D']), 10, base=10))
    cm = np.array([(0, 0, 0, 0)] + list(plt.cm.viridis_r(np.linspace(0, 1, len(zbin_cp)))))
    cmap_param = {'levels':zbin_cp, 'colors':cm}

    ax.set(ylim=(1.02, -.02))
    lab = ax.set_xlabel('Collective Generation', fontdict=fontdict)

    if title is not None:
        titl = ax.set_title(title, fontdict=fontdict)

    # Plot the contours...
    if zoom_first is None:
        im = ax.contourf(cp_density,
                         extent=[0, cp_density.shape[1], 0, 1],
                         **cmap_param)
    else:
        # Before the break
        im = ax_zoom.contourf(cp_density[:, :zoom_first],
                              extent=[0, zoom_first, 0, 1],
                              **cmap_param)
        # After the break.
        im = ax.contourf(cp_density[:, zoom_first:],
                         extent=[zoom_first, cp_density.shape[1], 0, 1],
                         **cmap_param)
        # Ajust ticks and labels positions
        ax_zoom.set(ylim=(1.02, -.02))
        ax.set(yticks=[])
        if title is not None:
            plt.setp(titl, position=(xcenter,plt.getp(titl, 'position')[1]))
        plt.setp(lab, position=(xcenter,plt.getp(lab, 'position')[1]))


    # Ticks
    atick = ax_zoom if ax_zoom is not None else ax
    ticks = atick.set(yticklabels=['Blue', 'Purple', 'Red'], yticks=[1, 0.5, 0])
    plt.setp(ticks[1][1], rotation=90, horizontalalignment='right', verticalalignment='center')
    atick.set_ylabel('Collective Colour', fontdict=fontdict)

    if colorbar:
        if cax is None:
            divider = make_axes_locatable(ax)
            cax = divider.append_axes("right", size="4%", pad=0.05)

        cbar = plt.gcf().colorbar(im,
                                  spacing='uniform',
                                  cax=cax,
                                  ticks=colorbar_ticks)

        cbar.ax.set_ylabel('Number of Collectives', fontdict=fontdict)
        cbar.ax.set_yticklabels(colorbar_ticks)


    return ax
```
