## Supplementary material for "Eco-evolutionary dynamics of nested Darwinian populations and the emergence of community-level heredity": Python code, figures and simulations: process.py

```
"""heredity/process.py -- extract data from evolution of heredity simulations.

This file is part of the ecological scaffolding package/ heredity model subpackage.
Copyright 2018 Guilhem Doulcier, Licence GNU GPL3+
--
"""

import logging
from collections import defaultdict
from functools import partial
import multiprocessing

import numpy as np
import pandas as pd
import estaudel.heredity.stochastic as model
import estaudel.heredity.deterministic as deter

logger = logging.getLogger('estaudel')
logger.setLevel(logging.DEBUG)
logger.propagate = False


def extract(out, pool=None, complete=True):
    """Populate the data attribute of an ecological scafolding Output object."""

    if pool is None:
        pool = multiprocessing.Pool(1)

    if 'cp_density' not in out.data or True:
        out.data['cp_density'], out.data['cp_value'] = extract_collective_phenotype(out)

    if 'individual_traits' not in out.data:
        individual_traits = extract_individual_traits(out)
        out.data['individual_traits'] = individual_traits

    if complete and 'individual_traits_density' not in out.data:
        individual_traits_density, individual_traits_bins = extract_individual_traits_density(
            out.data['individual_traits'])
        out.data['individual_traits_density'] = individual_traits_density
        out.data['individual_traits_bins'] = individual_traits_bins

    if complete and 'resident_id' not in out.data:
        out.data['resident_id'], out.data['resident_pheno'] = get_mab(out)

    if complete and 'resident_deter_traits' not in out.data:
        out.data['resident_deter_traits'] = pool.map(partial(deter.convert_phenotypes_to_lv, K=out.parameters['carrying_capacity']),
                                                     out.data['resident_pheno'])
    if complete and ('pstar' not in out.data or 'tstar' not in out.data):
        fixed_point_pheno = pool.map(
            deter.pstar, (A for _, A in out.data['resident_deter_traits']))
        out.data['pstar'] = np.vectorize(lambda x: (fixed_point_pheno[x]
                                                    if not np.isnan(x)
                                                    else np.nan))(out.data['resident_id'])

    if complete and 'tstar' not in out.data:

        tcrit_list = pool.starmap(partial(deter.tstar, B=out.parameters['B'], precise=True),
                                  out.data['resident_deter_traits'])
        out.data['tstar'] = np.vectorize(lambda x: (tcrit_list[int(x)]
                                                    if not np.isnan(x)
                                                    else np.nan))(out.data['resident_id'])


def extract_individual_traits(out, gen=None, colors=(0, 1), positions=(1, 3)):
    """Create a dict of tables with `value (of_trait), number (of_individuals), generation`"""
    traits = {}

    if gen is None:
        gen = out.current_gen
    for color in colors:
        for pos in positions:
            key = (model.COLOR_NAMES[color], model.POS_NAMES[pos])
            traits[key] = []
            for n in range(gen):
                traits[key].append(defaultdict(lambda: 0))
                for d in range(out.phenotype.shape[3]):
                    for ty in range(out.phenotype.shape[0]):
                        if out.phenotype[ty, 0, n, d] == color:
                            traits[key][-1][out.phenotype[ty, pos, n, d]
                                            ] += out.state[ty, n, d]
            col = []
            for g, x in enumerate(traits[key]):
                col.append(pd.DataFrame(pd.Series(x)).reset_index())
                col[-1].columns = ['value', 'number']
                col[-1]['generation'] = g
            traits[key] = pd.concat(col)
    return traits


def extract_collective_phenotype(out, nstep=100, gen=None):
    """
    Args:
        out (Output object) with phenotype, state and current_gen attributes.
        nstep (int): number of bins for the density
        gen (int): number of generations to process (if None, process all)
    Return:
        density (np.array): a (nstep, generation) array giving the histogram of type 1 individuals.
        trait (np.array): (generation, droplet) array containing the proportion of type 1 individuals.
    """
    if gen is None:
        gen = out.current_gen + 1
    trait = np.zeros((gen, out.state.shape[2]))
    density = np.zeros((nstep, gen))
    bins = np.linspace(0, 1, nstep+1)
    for n in range(gen):
        for d in range(out.state.shape[2]):
            trait[n, d] = np.sum(out.state[:, n, d]*out.phenotype[:,
                                                                  0, n, d])/out.state[:, n, d].sum()
        density[:, n] = np.histogram(trait[n, :], bins)[0]
        density[:, n] /= density[:, n].sum()
    return density, trait


def extract_individual_traits_density(individual_traits, nstep=100):
    bins = {}
    for key in frozenset([x[1] for x in individual_traits.keys()]):
        mx = np.max(individual_traits[(model.COLOR_NAMES[0], key)]['value'].values)
        mi = np.max(individual_traits[(model.COLOR_NAMES[0], key)]['value'].values)
        mx = max(mx, np.max(
            individual_traits[(model.COLOR_NAMES[1], key)]['value'].values))
        mi = min(mi, np.min(
            individual_traits[(model.COLOR_NAMES[1], key)]['value'].values))
        bins[key] = np.linspace(mi, mx, nstep+1)
        logger.info("Individual trait: {} -> {} {}".format(key, mi, mx))
    individual_traits_density = {}
    for key in individual_traits.keys():
        gen = individual_traits[key].generation.max()
        individual_traits_density[key] = np.zeros((nstep, gen+1))
        for n, df in individual_traits[key].groupby('generation'):
            individual_traits_density[key][:, n], _ = np.histogram(df['value'].values,
                                                                   weights=df['number'].values,
                                                                   bins=bins[key[1]])
    return individual_traits_density, bins


def get_mab(output, gen=None):
    """
    Get the resident phenotype in each collective.
    the form is resident[phenotype][generation][id_collective]
    """
    if gen is None:
        gen = output.current_gen
    resident = {}
    u = 0
    resident_identity = np.zeros((gen, output.state.shape[2]), dtype=int)
    for n in range(gen):
        for d in range(output.state.shape[2]):
            try:
                most_abundant = [None, None]
                for color in 0, 1:
                    mask = output.phenotype[:, 0, n, d] == color
                    most_abundant[color] = np.arange(output.state.shape[0])[
                        mask][np.argmax(output.state[mask, n, d])]
                key = tuple(map(tuple, output.phenotype[most_abundant, :, n, d]))
                if key not in resident:
                    resident[key] = int(u)
                    u += 1
                resident_identity[n, d] = resident[key]
            except ValueError:
                resident_identity[n, d] = np.nan
    resident, _ = zip(*sorted(resident.items(), key=lambda x: x[1]))
    return resident_identity, resident
```
