## Supplementary material for "Eco-evolutionary dynamics of nested Darwinian populations and the emergence of community-level heredity": Python code, figures and simulations: stochastic.py

```
""" heredity/stochastic.py -- Evolution of heredity thought experiment, stochastic model.
This file is part of the ecological scaffolding package/ heredity model subpackage.
Copyright 2018 Guilhem Doulcier, Licence GNU GPL3+
--

Phenotypes in this experiment have size 4:

[color, growth_rate, intra_type_interaction, inter_type_interaction]

- Color: c_i.
- Growth_rate: r_i .
- intra_type_interaction: a_i.
- inter_type_interaction: a'_i.

If N_i is the number of inidividuals of type i:

Birth rates are:
    b_i(N) = r_i*N_i
Death rate are:
    d_i(N) = r_i*N_i*[{sum_{j, st. c_i=c_j} N_j * a_j + {sum_{j, st. c_i!=c_j} N_j * a'_j]
"""
import numpy as np

COLOR_NAMES = ['red', 'blue']
POS_NAMES = ['color', 'growth_rate', 'interaction_intra', 'interaction_inter']

# Stochastic model


def bd_rates(phenotypes, state, K):
    """ Return the density dependent birth and death rates of the phenotypes.

    Used as argument for stochastic.bdm_process.

    phenotypes (array): Description of the types
    state (array of int): Number of individual of each type
    K (float): Carrying capacity
    """
    rates = np.zeros((len(state), 2))

    # Birth rates
    rates[:, 0] = np.multiply(state, phenotypes[:, 1])

    # Death rates
    competition = np.array([np.sum([N/K * phenotypes[j, 2+int(phenotypes[j, 0] != phenotypes[i, 0])]
                                    for j, N in enumerate(state)])
                            for i in range(len(state))])
    rates[:, 1] = np.multiply(np.multiply(state, competition), phenotypes[:, 1])
    return rates


def collective_fitness(phenotypes, state, var=1, goal=.5):
    """Return the collective fitness of a collective with `state[i]`
    individuals of type `phenotypes[i]`.

    Used in argument in escaffolding.collective_generation.
    """
    prop = np.sum([n for g, n in zip(phenotypes, state) if g[0] == 0]) / state.sum()
    return 1/(var*np.sqrt(np.pi*2)) * np.exp(-0.5*((goal-prop)/var)**2)


def gen_collective(max_types, B, green=(0, 1, .8, .3), red=(1, 1, .9, .3), proportion=None):
    """Return a (phenotypes, sate) tuple encoding a collective.

    - maxTypes (int): Maximum number of different phenotypes in this collective.
    - B (int): Initial population size.
    - green (array) and red (array): Initial trait values for red and green types.
    - proportion (float): initial proportion of red and green individuals.
                          If none, it will be taken at random.
    """

    phenotypes = np.zeros((max_types, 4))
    for i in range(max_types):
        phenotypes[i, :] = green if i % 2 == 0 else red

    state = np.zeros(max_types)
    if proportion is None:
        proportion = np.random.random()
    state[0] = int(B*proportion)
    state[1] = B - state[0]

    return phenotypes, state
```
