## Supplementary material for "Eco-evolutionary dynamics of nested Darwinian populations and the emergence of community-level heredity": Python code, figures and simulations: treeview.py

```
"""treeview - A small library dedicated to plot trees of the
collective genelaogy with cairo"""

import numpy as np
import cairo
from collections import Counter, defaultdict, deque
import math

def simple_namer(g,i):
    if g is None or i is None:
        return 'ROOT'
    return "G{:05}D{:03}".format(g,i)


# Extract the data from the Output object
def extract_tree(output, G=None):
    if G is None:
        G = output.parameters['N']-1
    cpval = make_node_info(simple_namer,  output.data['cp_value'])
    branches, nodes = make_tree(output.parents[:G], namer=simple_namer)
    generation, position = layout(branches, sortby=cpval)
    return {'branches':branches, 'nodes':nodes, 'position':position, "generation":generation, 'colour':cpval}

def layout(branches, sortby=None, root='ROOT', delroot=True):
    height = {root:0}
    width = {root:0}
    current = root
    if sortby is None:
        sortf = lambda x:1
    else:
        sortf = lambda k: sortby[k]
    to_visit = deque()
    width_at_height = defaultdict(lambda:0)
    while current is not None:
        children = sorted([b for b in branches if b[0]==current], key = lambda b:sortf(b[1]))
        for parent,child in children:
            to_visit.append(child)
            height[child] = height[parent] + 1
            width_at_height[height[child]] += 1
            width[child] = width_at_height[height[child]]
        try:
            current = to_visit.popleft()
        except IndexError:
            current = None
    if delroot:
        del height[root]
        del width[root]
    return height, width

def make_tree(parents, namer=simple_namer):
    """
    Args:
        parents (np.array): P[g,d] index of the parent of droplet 'd' at generation 'g'.
        namer (function) maps the (d,g) to an unique name.
    return:
        branches (list) # pairs of connected node.
        position (dict) # Un-tangled y position.
        generation (dict) # Generation of node.
    """
    try:
        G, N = parents.shape
    except AttributeError:
        parents = np.array(parents)
        G, N = parents.shape
        idx = np.arange(N)

    idx = np.arange(N)
    branches = [('', namer(None, None))]+[(namer(None, None), namer(0, i)) for i in idx]

    for g in range(1, G+1):
        pos = 0
        for i in idx:
            id_child = idx[parents[g-1] == i]
            for child in id_child:
                branches.append((namer(g-1, i), namer(g, child)))
                pos += 1

    childs = frozenset([b[1] for b in branches])
    parents = frozenset([b[0] for b in branches])
    nodes = parents.union(childs)
    return branches, nodes

def make_node_info(namer, data, info=None):
    if info is None:
        info = dict()
    G, N = data.shape
    for g in range(G):
        for i in range(N):
            info[namer(g,i)] = data[g,i]
    return info

def plot_ring_slice(ctx, cx,cy,r,angle_1,angle_2,shrink=0.8):
    r1, r2 = r, r*shrink
    ctx.move_to(cx+np.cos(angle_1)*r2,cy+np.sin(angle_1)*r2)
    ctx.line_to(cx+np.cos(angle_1)*r1,cy+np.sin(angle_1)*r1)
    ctx.arc(cx, cy, r1, angle_1, angle_2)
    ctx.move_to(cx+np.cos(angle_2)*r1,cy+np.sin(angle_2)*r1)
    ctx.line_to(cx+np.cos(angle_2)*r2,cy+np.sin(angle_2)*r2)
    ctx.arc_negative(cx, cy, r2, angle_2, angle_1)

def plot_ring_prop(ctx, cx, cy, r, prop, shrink=0.8, colors=(
((0x4e/255,0x79/255,0xa7/255)), (0xe1/255,0x57/255,0x59/255), ), alpha=1):
    ctx.set_source_rgba(*colors[0], alpha)
    plot_ring_slice(ctx, cx,cy,r,0, prop*2*math.pi,shrink=shrink)
    ctx.fill()
    ctx.set_source_rgba(*colors[1], alpha)
    plot_ring_slice(ctx, cx,cy,r, prop*2*math.pi, 2*math.pi,shrink=shrink)
    ctx.fill()
    ctx.set_source_rgb(1,1,1)
    ctx.arc(cx, cy, r*shrink, 0, 2*math.pi)
    ctx.fill()

def draw_tree_rings(branches, xinfo, yinfo, zinfo, oinfo=None, width=1024, height=600, size=6, label=None, line_width=3):
    # Draw tree
    childs = frozenset([b[1] for b in branches])
    parents = frozenset([b[0] for b in branches])
    nodes = parents.union(childs)

    xmin = np.min(np.array([*xinfo.values()]))
    xmax = np.max(np.array([*xinfo.values()]))
    ymin = np.min(np.array([*yinfo.values()]))
    ymax = np.max(np.array([*yinfo.values()]))
    leaves = [n for n,x in xinfo.items() if x==xmax]

    if oinfo is None:
        oinfo = {n:1 for n in nodes}
    elif oinfo == 'child':
        oinfo = {n:1 if n in parents or n in leaves else 0.3 for n in nodes}
    elif oinfo == 'coal':
        coal = coalescent(leaves, branches)
        oinfo = {n:1 if n in coal else 0.3 for n in nodes}

    xscale = lambda k: (width-3*size)*(xinfo[k]-xmin)/(xmax-xmin) + 1.5 * size
    yscale = lambda k: (height-3*size)*(yinfo[k]-ymin)/(ymax-ymin) + 1.5 * size
    oscale = lambda k: oinfo[k] if k in oinfo else 1
    zscale = lambda k: zinfo[k]

    surface = cairo.ImageSurface(cairo.FORMAT_ARGB32, width, height)
    ctx = cairo.Context(surface)

    ctx.scale(1, 1)  # Normalizing the canvas
    ctx.set_source_rgba(0.1, 0.1, 0.1, 1)  # Solid color
    ctx.set_line_width(line_width)

    for parent, child in branches:
        try:
            ctx.set_source_rgba(0.1, 0.1, 0.1, oscale(child))
            ctx.move_to(xscale(parent), yscale(parent))
            ctx.curve_to((xscale(child)+xscale(parent))/2, yscale(parent),
                         (xscale(parent)+xscale(child))/2, yscale(child),
                         xscale(child), yscale(child))
            ctx.stroke()
        except KeyError:
            pass

    for node in nodes:
        try:
            plot_ring_prop(ctx, xscale(node), yscale(node),
                           size, zscale(node),
                           shrink=0.5,
                           alpha=oscale(node))
        except KeyError:
            pass

    if label is not None:
        print(label)
        ctx.select_font_face("Arial", cairo.FONT_SLANT_NORMAL, 
                             cairo.FONT_WEIGHT_BOLD)
        ctx.set_font_size(200)
        ctx.set_source_rgba(0.1, 0.1, 0.1, 1)
        ctx.move_to(0.5,0.1)
        ctx.show_text(label*30)
        ctx.fill()
    return surface

def coalescent(nodes, branches):
    """Compute the coalescent of some nodes.

    Given a list of nodes and a list of branches, return the set of
    nodes in the coalescent tree of the input.
    """
    parent_map = {child:parent for parent,child in branches}
    coal = set(nodes)
    parents = set(nodes)
    while len(parents):
        parents = set([parent_map[n] for n in parents if n in parent_map])
        coal = coal.union(parents)
    return coal

def descent(nodes, branches):
    """Compute the descent of some nodes.
    Given a list of nodes and a list of branches, return the set 
    of nodes in the descent tree of the input.
    """
    desc = set(nodes)
    children = set(nodes)
    while len(children):
        children = set([child for parent, child in branches if parent in children])
        desc = desc.union(children)
    return desc
```
