## Supplementary material for "Eco-evolutionary dynamics of nested Darwinian populations and the emergence of community-level heredity": Python code, figures and simulations: stochastic.py

```
""" stochastic.py -- functions to perform stocastic birth-death-mutation process.
This file is part of the ecological scaffolding package.
Copyright 2018 Guilhem Doulcier, Licence GNU GPL3+
"""

from itertools import chain
import numpy as np

def discrete_bdm_process(T, steps, skip, state, phenotypes,
                         mutation_rate, mutation_function, rate_function):
    """ Perform a stochastic simulation of the birth-death process with mutations.

    Args:
        T (float): Final time. The process is simulated on [0,T].
        steps (int): number of time division.
        skip (int): only save one every `skip` state
        state (np.array of int): the number of individual of each type.
        phenotypes (np.array of floats): the value of the trait of each type.
        mutation_rate (float): Probability of mutation for a birth event.
        mutation_function (phenotype -> phenotype): Perform the mutation.
        rate_function (state,phenotypes->iter): Return growth rate of each type. Ecological model.

    Returns:
        trajectory (np.array of int, shape= (len(state), steps))
        pheno_traj (np.array of floats, shape= (len(state), steps))
    """
    M = len(state) # number of different phenotypes

    # Initialize the data structure for the trajectory
    trajectory = np.zeros((M, steps//skip), dtype=int)
    trajectory[:, 0] = state
    new_state = np.zeros(M)
    tlist = np.zeros(steps//skip)
    dt = np.diff(np.linspace(0,T,steps))[0]

    history = [{'first':0,'last':steps, 'pos':i,'phenotype':phenotypes[i, :].copy()} for i in range(len(state))]
    key_pheno = np.arange(len(state))
    mutid = len(state)

    for t in range(1, steps):
        # At each time step we compute the new rates
        rates = rate_function(phenotypes, state)
        nmutants = np.zeros(M, dtype=int)

        for i in range(M):
            if state[i]:
                new_state[i] = birth_death_tauleap(state[i], rates[i, 0], rates[i, 1], dt)
                birth = new_state[i] - state[i]
                if birth > 0:
                    # Each birth event may give rise to a potential
                    # mutation with probability mutation_rate.
                    nmutants[i] = np.random.binomial(birth, mutation_rate)


        # Treat the mutations.
        if mutation_rate and nmutants.sum():
            for color in (0, 1):
                # nmutant[i] contains how many potential mutations
                # arose during this time step for type i.  But we only
                # apply mutations if we have a type spot to store
                # them.

                # Create a list that contains the index of the phenotype of
                # every individual that mutated during the last time interval.
                # i.e. if nmutant = [1,3,2], parent_list = [0,1,1,1,2,2].
                parent_list = list(chain(*[[i]*(n if phenotypes[i, 0]==color else 0)
                                           for i, n in enumerate(nmutants)]))

                # List the empty phenotype slots, this is where we will put the mutants.
                #mutants_list = np.arange(M)[state == 0]
                mutants_list = np.arange(M)[np.logical_and(state == 0, phenotypes[:, 0] == color)]

                # If we have too many potential parents compared to the number of open slots
                # for phenotype, we select them uniformely at random.
                # Note: this is quite an important restriction of our simulation !
                if len(parent_list) > len(mutants_list):
                    parent_list = np.random.choice(parent_list,
                                                   size=len(mutants_list),
                                                   replace=False)

                # Go through the mutations and do the apply them.
                for pos_parent, pos_mutant in zip(parent_list, mutants_list):
                    phenotypes[pos_mutant, :] = mutation_function(phenotypes[pos_parent, :])
                    new_state[pos_mutant] = 1
                    new_state[pos_parent] -= 1

                    history[key_pheno[pos_mutant]]['last'] = t
                    key_pheno[pos_mutant] = mutid
                    mutid += 1
                    history.append({'first':t,
                                    'last':steps,
                                    'pos':pos_mutant,
                                    'phenotype':phenotypes[pos_mutant, :].copy()})

        # We copy the content of new_state in state.
        state[:] = new_state[:]
        # We save once in a while.
        if t%skip == 0:
            trajectory[:, t//skip] = state
            tlist[t//skip] = t*dt
    return {'state':state,
            'phenotype': phenotypes,
            'trajectory':trajectory,
            'times':tlist,
            'history':history}

def normal_mutation_abs(old, effect):
    """ Mutation function
    A trait is selected at random and mutated by adding a normal random variable.
    The new trait is reflected around 0 (absolute value).

    Args:
       old (np.array): old phenotype
       effect (dict): for each mutable position: position : standard deviation.

    Used as argument for stochastic.bdm_process
    """
    pos = int(np.random.choice(list(effect.keys())))
    new = old.copy()
    new[pos] = abs(np.random.normal(old[pos], effect[pos]))
    return new


def birth_death_tauleap(n: int, b: float, d: float, dt: float):
    """Return a realization of the discretisation of a birth-death process.

    Args:
    - n (int): initial number of individuals.
    - b (float): birth rate
    - d (float): death rate.
    - dt (float): tau.

    Return the new number of individual.

    This ensure that the new population size is not negative by
    returning 0 at maximum.
    """
    return max(n + np.random.poisson(b*dt) - np.random.poisson(d*dt), 0)


##### EXPERIMENTAL CODE BELOW #####

def get_p0_rho(b: float, d: float, dt: float):
    """ Return the probabilities for a birth-death process with:

    Args:
        b (float): birth rate.
        d (float): death rate.
        dt (float): time of observation.

    Return the tuple (p0,rho)

    p0 is the probability that the process is extinct at after dt.
    rho is such that P(Z_t=k | non_extinction ) = (1-rho)^k rho.
    """
    if b-d < 0:
        e = np.exp((b-d)*dt)
        p0 = d * (1 - e) / (d-b*e)
        rho = (d-b) / (d-b*e)
    if b==d:
        p0 = b*dt / (1 + b*dt)
        rho = 1 / (1 + b*dt)
    else:
        e = np.exp(-(b-d)*dt)
        p0 = d * (1 - e) / (b-d*e)
        rho = (b-d)*e / (b-d*e)
    return p0, rho

def birth_death_discrete(n: int, b: float, d: float, dt: float):
    """Return a realization of the discretisation of a birth-death process.

    Args:
    - n (int): initial number of individual.
    - b (float): birth rate
    - d (float): death rate.
    - dt (float): time of observation.

    /!\ FIXME: Timescales are wrong !!!! /!\

    Return the new number of individual.
    """
    p0, rho = get_p0_rho(b, d, dt)

    # number of individual whose decent is not extinct after dt.
    not_extinct = np.random.binomial(n, 1-p0)

    # new number of individuals
    new_n = np.sum(np.random.geometric(rho, size=not_extinct))
    return new_n
```
