## Supplementary material for "Eco-evolutionary dynamics of nested Darwinian populations and the emergence of community-level heredity": Python code, figures and simulations: config.py

```
from matplotlib.colors import ListedColormap, LinearSegmentedColormap
import os

PANEL_DICT = dict(xy=(1, 1), xycoords="axes fraction",
                 ha='center', va='center',
                 bbox=dict(boxstyle="round",
                           ec=(.1, 0.1, .1),
                           fc=(1., 1, 1),))

 
FIGPATH = 'fig'
SUPFIGPATH = 'supfig'

for path in [FIGPATH,SUPFIGPATH]:
    if not os.path.exists(path):
        os.mkdir(path)

FONT_DICT  ={'fontweight': 'normal', 'size':'x-large'}


cdict1 = {'red':   ((0, 207/256, 207/256),
                    (1.0, 0.0, 0.0)),

         'green': ((0.0, 0.0, 0.0),
                   (1, 171/256, 171/256)),

         'blue':  ((0, 54/256, 54/256),
                   (1, 212/256, 212/256))
         }

cdict = {'red':   ((1.0, 207/256, 207/256),
                    (0, 0.0, 0.0)),

         'green': ((1, 0.0, 0.0),
                   (0, 171/256, 171/256)),

         'blue':  ((1, 54/256, 54/256),
                   (0, 212/256, 212/256))
         }

BLUE_RED_r = LinearSegmentedColormap('BlueRed_r', cdict1)
BLUE_RED = LinearSegmentedColormap('BlueRed', {k:v[::-1] for k,v in cdict.items()})
```
