## Supplementary material for "Eco-evolutionary dynamics of nested Darwinian populations and the emergence of community-level heredity": Python code, figures and simulations: constrained_trajectories.ipynb

````
# %% [markdown]
# Constrained trajectories

This figure illustrate what happen if only one of the two pairs of traits (growth rate, or inter-color interaction rate) that were free to evolve become fixed.  

Now we can look at a trajectory in a 2D trait-space ! 

# %% [code]
from functools import partial
import numpy as np
import matplotlib.pyplot as plt
from mpl_toolkits.axes_grid1 import make_axes_locatable
from matplotlib.colors import ListedColormap, LinearSegmentedColormap

import numpy as np
import estaudel.escaffolding as escaffolding
import estaudel.heredity.process
from collections import Counter
import estaudel.heredity.deterministic

import os 
from config import PANEL_DICT, FIGPATH, BLUE_RED

# %% [markdown]
# Data

Here are the commands to generate this data

```bash

# Mutations only on the inter-color interaction rate (pos 3)
python3 ./full_trajectory.py --N 300 --mutation_effect {3:0.05} --initial_type0 "(0,25,0.8,0.7)" --initial_type1 "(1, 25, 0.7, 0.15)"

# Mutations only on the growth rate (pos 1)
python3 ./full_trajectory.py --N 300 --carrying_capacity 1e20 --mutation_effect {1:0.05}```


# %% [code]
path_a = "data/constrained/20180629_200200_initial_type1(1, 25, 0.7, 0.15)_mutation_effect{3:0.05}_N300_initial_type0(0,25,0.8,0.7).pkle"
path_r = "data/constrained/20180629_200142_mutation_effect{1:0.05}_carrying_capacity1e20_N300.pkle"

# %% [code]
out_a = escaffolding.load(path_a)
print(out_a)
print("\n".join(["{}: {}".format(k,v) for k,v in sorted(out_a.parameters.items())]))

# %% [code]
out_r = escaffolding.load(path_r)
#out_r.data['tstar'] = None
#estaudel.heredity.process.extract(out_r)
#print(out_r)
print("\n".join(["{}: {}".format(k,v) for k,v in sorted(out_r.parameters.items())]))

# %% [markdown]
# Functions

# %% [code]
def get_resident_cloud(out, pos):
    x,y,z = [],[],[]
    for k,v in Counter(list(out.data['resident_id'].flatten())).items():
            x.append(out.data['resident_pheno'][k][0][pos])
            y.append(out.data['resident_pheno'][k][1][pos])
            z.append(v)
    mean_x, mean_y = [], []
    
    for gen in out.data['resident_id']:
        mean_x.append(np.sum([out.data['resident_pheno'][k][0][pos] * v for k,v in Counter(gen).items()])/len(gen))
        mean_y.append(np.sum([out.data['resident_pheno'][k][1][pos] * v for k,v in Counter(gen).items()])/len(gen))
    return x,y,z,mean_x, mean_y

# %% [markdown]
# Plots

# %% [code]
r_cloud = get_resident_cloud(out_r,1)
a_cloud = get_resident_cloud(out_a,3)

# %% [code]
r,A = estaudel.heredity.deterministic.convert_phenotypes_to_lv(
    np.array([out_a.parameters['initial_type0'], out_a.parameters['initial_type1']]), K=1)
xx = np.linspace(0,0.85)
Xa,Ya = np.meshgrid(xx,xx)
f = np.vectorize(lambda x,y: estaudel.heredity.deterministic.pstar(np.array([[A[0,0],y],[x,A[1,1]]])))
f = np.vectorize(lambda x,y : estaudel.heredity.deterministic.get_gfunc(r=[25,25],
                                                                        A=np.array([[A[0,0],y],[x,A[1,1]]]),
                 B=15)(0.5,T=1))

Za = f(Xa,Ya)

xx = np.linspace(np.min(np.array(r_cloud[:2])),np.max(np.array(r_cloud[:2])))
Xr,Yr = np.meshgrid(xx,xx)
f = np.vectorize(lambda x,y : estaudel.heredity.deterministic.get_gfunc(r=[x,y],A=None,B=None)(0.5,T=1))
Zr = f(Xr,Yr)

# %% [code]
scatter = True
plt.rc('font', size=15)
fig, ax = plt.subplots(1,2,figsize=(14,7))
ax = ax[::-1]
cax = make_axes_locatable(ax[0]).append_axes("right", size="4%", pad=0.11) 

PANEL_DICT.update({'xy':(0.05,0.95), 'size':20})
ax[1].annotate('A', **PANEL_DICT)
ax[0].annotate('B', **PANEL_DICT)


ticks = [0,0.25,0.5,0.75,1]
levels = np.linspace(0,1,20)

########### Mutating A #######################
# Background 
im = ax[0].contourf(Xa, Ya, Za, cmap=BLUE_RED, levels=levels)
ax[0].contourf(Xa, Ya, np.int_(np.isnan(Za)), colors='white', levels=[0.1,1],  hatches='/') 
ax[0].contour(Xa,Ya,Za,colors='w',linestyles=':',levels=[0.5])

# Colorbar
#fig.colorbar(im, cax=cax[0], ticks=ticks,
#             label=r'Stable fixed point of $G$ when $T \to \infty$, $p^*$')
#cax[0].hlines(0.5,0,1,color='C0',linestyles=':')

# Trajectory
if scatter:
    ax[0].scatter(a_cloud[0], a_cloud[1], s=a_cloud[2], marker='.', alpha=.1, color='k', label='Resident phenotype')
ax[0].plot(a_cloud[3],a_cloud[4],  lw=3, color='w',  label='Mean resident phenotype')
ax[0].plot(a_cloud[3],a_cloud[4], '>', markevery=(100,100), 
            markersize=12,
           lw=3, color='w',  label='Mean resident phenotype')

# Set axis
ax[0].set(
        xlabel=r'Inter-color interaction rate of type 0 resident', 
        ylabel=r'Inter-color interaction rate of type 1 resident',
        title='Fixed growth rates',
        xlim=(Xa[0,0],Xa[-1,-1]),
        ylim=(Ya[0,0],Ya[-1,-1]),
    xticks=[0,0.2,0.4,0.6,0.8],
      yticks=[0,0.2,0.4,0.6,0.8],
      yticklabels=[0,0.2,0.4,0.6,0.8],
      xticklabels=[0,0.2,0.4,0.6,0.8]
)
########### Mutating r #########################

# Background image
im = ax[1].contourf(Xr,Yr,Zr,cmap=BLUE_RED,levels=levels)
ax[1].contour(Xr,Yr,Zr,colors='w',levels=[0.5], linestyles=':')

# Trajectory
if scatter:
    ax[1].scatter(r_cloud[0],r_cloud[1], s=r_cloud[2], marker='.', alpha=.1, color='k')
ax[1].plot(r_cloud[3],r_cloud[4], lw=3, color='w', label='Mean resident phenotype')
ax[1].plot(r_cloud[3],r_cloud[4], ">", markevery=(200,200),
           markersize=12, lw=1, color='w',
           label='Mean resident phenotype')

# Colorbar
fig.colorbar(im,cax=cax, label="G(0.5)",ticks=ticks)
cax.hlines(0.5,0,1,color='w', linestyles=':')
cax.set_yticklabels(ticks)
ax[1].set(
    title='Fixed interaction rates',
    xlabel='Growth rate of the type 0 resident', 
    ylabel='Growth rate of the type 1 resident',
    xlim=(Xr[0,0],Xr[-1,-1]),
    ylim=(Yr[0,0],Yr[-1,-1]))
fig.tight_layout()
fig.savefig(os.path.join(FIGPATH,'constrained_trajectories.png'), bbox_inches='tight')

# %% [markdown]
## Alternative fig


# %% [code]
fig, ax = plt.subplots(1,1,figsize=(10,10))
ax = [ax]
cax = [make_axes_locatable(a).append_axes("right", size="4%", pad=0.11) for a in ax]
#ax[0].annotate('B', **panel_dict)


ticks = [0,0.25,0.5,0.75,1]
levels = np.linspace(0,1,20)

########### Mutating A #######################
# Background 
im = ax[0].contourf(Xa, Ya, Za, cmap=blue_red1, levels=levels)
ax[0].contourf(Xa, Ya, np.int_(np.isnan(Za)), colors='white', levels=[0.1,1],  hatches='/') 
ax[0].contour(Xa,Ya,Za,colors='w',linestyles=':',levels=[0.5])

# Colorbar
fig.colorbar(im, cax=cax[0], ticks=ticks,
             label=r'Stable fixed point of $G$ when $T \to \infty$, $p^*$')
cax[0].hlines(0.5,0,1,color='C0',linestyles=':')

# Trajectory
#ax[0].scatter(a_cloud[0], a_cloud[1], s=a_cloud[2], marker='.', alpha=.1, color='k', label='Resident phenotype')
ax[0].plot(a_cloud[3],a_cloud[4],  lw=3, color='w',  label='Mean resident phenotype')
ax[0].plot(a_cloud[3],a_cloud[4], '>', markevery=12, lw=3, color='w',  label='Mean resident phenotype')

# Set axis
ax[0].set(
        xlabel=r'Inter-color interaction rate of type 0 resident, $a_0^{inter} \times K$', 
        ylabel=r'Inter-color interaction rate of type 1 resident, $a_1^{inter} \times K$',
        title='Fixed growth rates, mutating interaction rates',
        xlim=(Xa[0,0],Xa[-1,-1]),
        ylim=(Ya[0,0],Ya[-1,-1])
)
#plt.savefig('/home/guilhem/phd/www/slides/190326_IBENS/slides/bifA.svg')

# %% [code]
fig, ax = plt.subplots(1,1,figsize=(7,7))
ax = [ax, None]
cax = [make_axes_locatable(a).append_axes("right", size="4%", pad=0.11) for a in ax if a]
ticks = [0,0.25,0.5,0.75,1]
levels = np.linspace(0,1,20)

########### Mutating A #######################
# Background 
im = ax[0].contourf(Xa, Ya, Za, cmap=BLUE_RED, levels=levels)
ax[0].contourf(Xa, Ya, np.int_(np.isnan(Za)), colors='white', levels=[0.1,1],  hatches='/') 
ax[0].contour(Xa,Ya,Za,colors='w',linestyles=':',levels=[0.5])

# Colorbar
fig.colorbar(im, cax=cax[0], ticks=ticks,
             label=r'Stable fixed point of $G$ when $T \to \infty$, $p^*$')
cax[0].hlines(0.5,0,1,color='C0',linestyles=':')

# Trajectory
ax[0].scatter(a_cloud[0], a_cloud[1], s=a_cloud[2], marker='.', alpha=.1, color='k', label='Resident phenotype')
ax[0].plot(a_cloud[3],a_cloud[4],  lw=3, color='w',  label='Mean resident phenotype')
ax[0].plot(a_cloud[3],a_cloud[4], '>', markevery=12, lw=3, color='w',  label='Mean resident phenotype')

# Set axis
ax[0].set(
        xlabel=r'Inter-color interaction rate of type 0 resident, $a_0^{inter} \times K$', 
        ylabel=r'Inter-color interaction rate of type 1 resident, $a_1^{inter} \times K$',
        title='Fixed growth rates, mutating interaction rates',
        xlim=(Xa[0,0],Xa[-1,-1]),
        ylim=(Ya[0,0],Ya[-1,-1])
)

#fig.savefig(os.path.join(FIGPATH,'mutA.png'), bbox_inches='tight', dpi=300)
#fig.savefig(os.path.join(FIGPATH,'mutA.svg'), bbox_inches='tight', dpi=600)
#!ls -a fig/mutA.png

# %% [code]
````
