## Supplementary material for "Eco-evolutionary dynamics of nested Darwinian populations and the emergence of community-level heredity": Python code, figures and simulations: fixed_point_as_function_of_B_T.ipynb

```
# %% [markdown]
# Effect of $B,T$ on $G$

# %% [markdown]
In this figure we show the position and stability of the fixed points of $G$ as a function of the duration of the growth phase $T$ a d Bottlneck size $B$.

# %% [code]
from collections import OrderedDict
import numpy as np 
import matplotlib.pyplot as plt
import estaudel.heredity.deterministic as ehd
from mpl_toolkits.axes_grid1 import make_axes_locatable

%matplotlib inline
import os 
from config import FIGPATH, PANEL_DICT
from config import PANEL_DICT, FIGPATH, SUPFIGPATH, BLUE_RED
import matplotlib.gridspec as gridspec
import matplotlib.pyplot as plt

# %% [markdown]
# Parameters

# %% [code]
r = np.array([1.2,1])*30
Aset = {
 'coex': np.array([[0.7, 0.3],
                 [0.2, 0.6]]),
 '1excludes0':np.array([[0.6, 0.7],
                      [0.4, 0.3]]),
 '1excludes0_2':np.array([[0.6, 1.2],
                      [0.1, 0.4]]),
 '0excludes1':np.array([[0.3, 0.6],
                       [0.5, 0.8]]),
 'bistable':np.array([[0.3, 0.7],
                      [0.8, 0.2]]),
}
Alist = [ '0excludes1',  '1excludes0_2', 'bistable', 'coex']
Anames = {
 'coex': "Asymptotic Coexistance,\n $a_{01} < a_{11}$ and $a_{10} < a_{00}$",
 '1excludes0_2': "Competitive exclusion of the fast grower,\n $a_{11} < a_{01}$ and $a_{10} < a_{00}$",
 '0excludes1': "Competitive exclusion of the slow grower,\n $a_{01} < a_{11}$ and $a_{00} < a_{10}$",
 'bistable': "Asymptotic Bistability,\n $a_{11} < a_{01}$ and $a_{00} < a_{10}$",
}
Anames = {
 'coex': "Asymptotic Coexistence",
 '1excludes0_2': "Competitive exclusion of the fast grower",
 '0excludes1': "Competitive exclusion of the slow grower",
 'bistable': "Asymptotic Bistability",
}
ehd.pstar(Aset['coex'])

# %% [code]
def bstar(t, side, r, A):
    x,y = (0,1) if side==0 else (1,0)
    alpha = r[x]*r[y]*(A[x,x]-A[y,x])/(r[x]*A[x,x]-r[y]*A[y,x])
    return  -(np.exp(-r[x]*t) - np.exp(-alpha*t)) / (A[x,x]*(1 - np.exp(-alpha*t)))

# %% [code]
x = np.linspace(0,1,300)

# %% [code]
lspace = np.logspace(-5, 1, 100)
tspace = np.linspace(0, 1, 100)
X,Y = np.meshgrid(tspace,lspace)
Z = np.zeros_like(X) + np.nan
for i,B in enumerate(lspace):
    asymptotic_pstar = ehd.pstar(Aset['coex'])
    g = ehd.get_gfunc(r,Aset['coex'],B)
    dd = ehd.continuation_on_T(g, t0=1, tf=1e-3, start=asymptotic_pstar)
    Z[i,:] = dd[1][::-1]
Z[np.abs(Z-1)<1e-5] = np.nan

# %% [code]
def plot_tb(r,A, tspan=np.linspace(0, 1, 100), betamin=1e-2, ax=None, colored_line=True):
    line = [[],[]]
    amax = A.max()
    fast_grower, slow_grower = int(r[1]>r[0]), int(r[1]<r[0])
    color = {0: BLUE_RED(1.0), 1:BLUE_RED(0.0)}
    cname = {0:'red',1:'blue'}

    for side in (0,1):
        line[side] = bstar(tspan, side=side, r=r,A=A)
        line[side][line[side]<0] = np.nan
        line[side][0] = line[side][1]
        
    if ax is None:
        ax = plt.gca()
    
    for side in (0,1):
        if not np.isnan(line[side][1]):
            c = {0:'r',1:'b'}
            ax.plot(tspan,line[side], color=c[side] if colored_line else 'k', 
                    lw=4)
    
    if True:
        alpha=1
        lowline = np.nanmin(np.vstack(line), 0)
        highline = np.nanmax(np.vstack(line), 0)
        if np.isnan(line[0][1]) and np.isnan(line[1][1]):
            ax.fill_between(tspan,0,1/amax, color=color[fast_grower], alpha=alpha,
                            label=cname[fast_grower].capitalize()+' stable')
        else:
            ax.fill_between(tspan,0,lowline, color=color[fast_grower], alpha=alpha,
                            label=cname[fast_grower].capitalize()+' stable')
        if not np.isnan(line[0][1]) and not np.isnan(line[1][1]):
            ax.fill_between(tspan,highline,1, color=color[slow_grower], alpha=alpha, 
                            label=cname[slow_grower].capitalize()+' stable')
            fast_is_low = lowline==line[fast_grower]
            fast_is_high = highline==line[fast_grower]
            if any(fast_is_low):
                ax.fill_between(tspan[fast_is_low],
                                line[fast_grower][fast_is_low],
                                line[slow_grower][fast_is_low],
                                color='purple', alpha=alpha, label='Purple stable')
            if any(fast_is_high):
                ax.fill_between(tspan[fast_is_high],
                                line[fast_grower][fast_is_high],
                                line[slow_grower][fast_is_high],
                                color='grey', alpha=alpha, label='Red and Blue stables')
        if np.isnan(line[fast_grower][1]) and not np.isnan(line[slow_grower][1]):
            ax.fill_between(tspan,highline,1, color='#bab0ac', alpha=alpha, label='Red and Blue stables')
    ax.semilogy()

    ax.set(ylim=(betamin*1/amax, 1/amax),
           xlim=(tspan.min(),tspan.max()))
    ax.set(ylabel='Bottleneck size, $B$',
           xlabel='Duration of growth phase, $T$')
    #ax.legend(loc='center right')

# %% [code]


# %% [code]
def plot_g(g, ax):
    gf = ehd.get_gfunc(r, Aset[g['Akey']], T=g['T'],B=g['B'] )
    l = ax.plot(x, [gf(xx) for xx in x], color='C1', lw=3)
    ax.set(xticks=[0,.5,1],yticks=[0,.5,1],xticklabels=[0,'',1],
           yticklabels=[0,'',1],xlabel='Newborn colour', ylabel='Adult colour')
    ax.plot([0,1],[0,1], color='k', alpha=0.1)
    #ax.set_title(g['name'])
    ax.grid()

# %% [code]
gfunc = []
gfunc.append({'B':1e-4, 'T':0.35, 'Akey':'0excludes1', 'color':BLUE_RED(1.0), 'name':'Fast Grower Fixation'})
gfunc.append({'B':1e-4, 'T':0.4, 'Akey':'1excludes0_2', 'color':BLUE_RED(0.0),'name':'Slow Grower Fixation' })
gfunc.append({'B':1e-4, 'T':.6, 'Akey':'bistable', 'color':'grey', 'name':'Unstable Purple Colour'}) 
gfunc.append({'B':1e-4, 'T':0.6, 'Akey':'coex', 'color':'purple', 'name':'Stable Purple Colour'}) 

# %% [code]
plt.rc('font', size=15)
fig,ax = plt.subplots(1,1,figsize=(6,6))

plot_tb(r, Aset['coex'], ax=ax, betamin=1e-5, colored_line=False)
ax.set(title=Anames['coex'])
im = ax.contourf(X,Y,Z, cmap=BLUE_RED, levels=np.linspace(0,1,30))

divider = make_axes_locatable(ax)
cax1 = divider.append_axes("right", size="5%", pad=0.05)
fig.colorbar(im, cax=cax1, ticks=[0,0.5,1])
cax1.set_yticklabels([0,.5,1]) 
cax1.set_ylabel('Fixed Point of G') 


ax.set(title='')
plt.rc('axes', labelcolor='w')
plt.rc('xtick', color='w', labelsize='small')
plt.rc('ytick', color='w', labelsize='small')
axin1 = ax.inset_axes([0.65, 0.65, 0.30, 0.30])
plot_g(gfunc[3], axin1)
axin2 = ax.inset_axes([0.08, 0.08, 0.20, 0.20])
plot_g(gfunc[0], axin2)
axin2.set(xlabel='',ylabel='')
plt.rc('axes', labelcolor='k')
plt.rc('xtick', color='k',  labelsize='medium')
plt.rc('ytick', color='k',  labelsize='medium')
plt.rc('font', size=15)
ax.set(xticks=[0,0.5,1], xticklabels=[0,0.5,1])
ax.set_ylim(1e-5,1)
ax.set_xlim(0,1)
plt.savefig(os.path.join(FIGPATH,'bifurcation_diagram.pdf'), bbox_inches='tight')

# %% [code]
grid_phase = gridspec.GridSpec(nrows=2, ncols=2, hspace=.1, wspace=.1, left=0.25, right=0.98)
grid_g = gridspec.GridSpec(nrows=4, ncols=1, left=0, right=0.18, hspace=.3)
grid_cax = gridspec.GridSpec(nrows=1, ncols=1, left=0.99, right=1)

plt.rc('font', size=15)

fig = plt.figure(figsize=(15,14))
axes_ph = [fig.add_subplot(x) for x in grid_phase]
axes_g = [fig.add_subplot(x) for x in grid_g]
cax = fig.add_subplot(grid_cax[0]) 
plt.tight_layout()

for Akey,ax in zip(Alist, axes_ph):
    plot_tb(r, Aset[Akey], ax=ax, betamin=1e-5)
    ax.set(title=Anames[Akey])
        
    if Akey == "coex":
        im = ax.contourf(X,Y,Z, cmap=BLUE_RED, levels=np.linspace(0,1,30))
    ax.set_ylim(1e-5,1)
    ax.set_xlim(0,1)


fd = {'fontweight': 'normal', 'size':'xx-large', 'verticalalignment':'center', 'horizontalalignment':'center'}
axes_ph[0].text( 0.5, 5e-3,'(1)', color='w', fontdict=fd)
axes_ph[1].text( 0.15, 1e-4,'(1)', color='w', fontdict=fd)
axes_ph[2].text( 0.15, 1e-4,'(1)', color='w', fontdict=fd)
axes_ph[3].text( 0.15, 1e-4,'(1)', color='w', fontdict=fd)
axes_ph[1].text( 0.072, 1e-1,'(3)', color='w', fontdict=fd)
axes_ph[1].text( 0.38, 5e-5,'(4)', color='w', fontdict=fd)

axes_ph[1].text( 0.65, 5e-3,'(2)', color='w', fontdict=fd)
axes_ph[2].text( 0.65, 5e-3,'(3)', color='w', fontdict=fd)
axes_ph[3].text( 0.65, 5e-3,'(4)', color='w', fontdict=fd)


for g, ax in zip(gfunc, axes_g):
    gf = ehd.get_gfunc(r, Aset[g['Akey']], T=g['T'],B=g['B'] )
    l = ax.plot(x, [gf(xx) for xx in x], color=g['color'], lw=3)
    ax.set(xticks=[0,.5,1],xticklabels=[0,.5,1],yticklabels=[0,.5,1],yticks=[0,.5,1], xlabel='Newborn colour', ylabel='Adult colour')
    ax.plot([0,1],[0,1], color='k', alpha=0.1)
    ax.set_title(g['name'])
    ax.grid()


axes_ph[1].set(ylabel='',yticklabels=[])
axes_ph[3].set(ylabel='',yticklabels=[])
axes_ph[0].set(xlabel='',xticklabels=[])
axes_ph[1].set(xlabel='',xticklabels=[])
for i in range(3):
    axes_g[i].set(xlabel='', xticklabels=[])

PANEL_DICT.update({'xy':(0.9,0.1), 'size':20})

for n,ax in zip('ABCD',axes_ph):
    ax.annotate(n, **PANEL_DICT)
for n,ax in zip('1234',axes_g):
    ax.annotate('{}'.format(n), **PANEL_DICT)
    
cbar = fig.colorbar(im, ax=axes_ph[-1], cax=cax, label='Stable fixed point of G')
cbar.set_ticks([1,0.5,0])
cbar.set_ticklabels(['Red','Purple','Blue'])
plt.savefig(os.path.join(SUPFIGPATH,'all_g_functions.pdf'), bbox_inches='tight')
```
