## Supplementary material for "Eco-evolutionary dynamics of nested Darwinian populations and the emergence of community-level heredity": Python code, figures and simulations: full_trajectory.ipynb

````
# %% [markdown]
# Full evolutionary trajectory

# %% [code]
from functools import partial
import numpy as np
import matplotlib.pyplot as plt
from mpl_toolkits.axes_grid1 import make_axes_locatable
from matplotlib.colors import ListedColormap
from collections import defaultdict
import matplotlib.gridspec as gridspec
import numpy as np

import estaudel.escaffolding as escaffolding
import estaudel.stochastic as stochastic
import estaudel.heredity.process as ehp
import estaudel.heredity.deterministic as ehd
import estaudel.heredity.stochastic as ehs

import os 
from config import FIGPATH, PANEL_DICT

# %% [markdown]
# Data

# %% [markdown]
These dataset are not included on the git repo (they are heavy) but they can be generated using the script `full_trajectory.py`

I.e. :
```bash
full_trajectory.py --selection "neutral"
full_trajectory.py --selection "rank"
```

# %% [code]
path_selection = "data/full_trajectory/full_trajectory.321.1_20180629_163801_selection\"rank\"_NPROC8.pkle"
path_neutral = "data/full_trajectory/full_trajectory.321.5_20180629_163801_NPROC8_selection\"neutral\".pkle"
prefix_out = "one"

data = (escaffolding.load(path_neutral), escaffolding.load(path_selection))
for d in data:
    print(d)
    print("\n".join(["{}: {}".format(k,v) for k,v in sorted(d.parameters.items())]))
    print('-'*80)

# %% [markdown]
# Individual and collectvie traits

# %% [code]
# Show a more detailed plot for the first Nbreak generations
Nbreak = 30

# %% [code]
# -------Axes limits-------
# Compute the maximal value of each trait, for axes limits. 
mx_trait = {}
for k in ('growth_rate','interaction_inter'):
    mx_trait[k] = (np.min([d.data['individual_traits_bins'][k].min() for d in data]),
                 np.max([d.data['individual_traits_bins'][k].max() for d in data]))

# Compute the maximum density and number of collectives 
mx = 0
D = 0
for d in data:
    D = max(D, d.parameters['D'])
    for k in d.data['individual_traits_density'].keys():
            mx = max(mx,d.data['individual_traits_density'][k].max())

# Bins for the colormap. 
rounder = np.vectorize(lambda x : round(x, -int(np.floor(np.log10(abs(x))))))

zbin = np.int_(rounder(list(np.logspace(0,np.log10(mx),8,base=10)[3:])))
zticks = zbin[::2]
zbin_names = ['{}k'.format(int(x)//1000) if x>1000 else x for x in zticks]
zbin_names = [0, r'$3{\times10^4}$', r'$2{\times}10^6$' ]


zbin_cp = [0.0000001]+list(np.logspace(0,np.log10(D),20,base=10))
zbin_cp = [-100,1]+list(np.logspace(1,np.log10(data[0].parameters['D']),25,base=10))
cmap_cp = np.array([(0,0,0,0)] + list(plt.cm.viridis_r(np.linspace(0,1,len(zbin_cp)))))

# %% [code]
mx = 0
D = 0
for i in [0,1]:
    D = max(D, data[i].parameters['D'])
    for pos in (1,3):
        for color in (0,1):
            mx = max(mx,data[i].data['individual_traits_density'][(ehs.COLOR_NAMES[color],
                                                                   ehs.POS_NAMES[pos])].max())

# %% [code]
plt.rc('font', size=15)
fig = plt.figure(figsize=(14,12))
tk =  [1,100,1000]
labels = ['ACE','BDF']

#-------- Set the axis structure ---- ###
gs = [gridspec.GridSpec(nrows=3, ncols=5, left=0.05, right=0.48,
                        wspace=0.01, hspace=0.1),
      gridspec.GridSpec(nrows=3, ncols=5, left=0.51, right=0.96,
                         wspace=0.01, hspace=0.1)]
g = gridspec.GridSpec(nrows=3, ncols=1, left=0.97, right=0.99,
                 wspace=0.01, hspace=0.1)

cax = []
cax.append(fig.add_subplot(g[0, 0]))
cax.append(fig.add_subplot(g[1, 0]))
cax.append(fig.add_subplot(g[2, 0]))
# ---------#

xcenter = 0.41
title = ['Neutral Regime', 'Selective Regime']
for i in (0,1):

    ax = []
    ax.append(fig.add_subplot(gs[i][0, 0]))
    ax.append(fig.add_subplot(gs[i][0, 1:]))
    
    ax.append(fig.add_subplot(gs[i][1, 0]))
    ax.append(fig.add_subplot(gs[i][1, 1:]))
    
    ax.append(fig.add_subplot(gs[i][2, 0]))
    ax.append(fig.add_subplot(gs[i][2, 1:]))
    ylabel_font = fontdict={'fontweight': 'normal', 'size':'x-large'}

    for j,x in enumerate([1,3,5]):
        if j==2:
            lab = ax[x].set_xlabel('Collective generation',)
            plt.setp(lab, position=(xcenter,plt.getp(lab, 'position')[1]))
        ax[x].set(yticks=[],
                  xticks=[5000,10000-1],
                  xticklabels=[r'$5{\times}10^3$','$10^4$'] if j==2 else [])
        ax[x].annotate(labels[i][j], **PANEL_DICT)
    for j,x in enumerate([0,2,4]):
        ax[x].set(xticks=[0,5,Nbreak],
                 xticklabels=[0,5,Nbreak] if j==2 else [])

    
    ticks = ax[0].set(ylim=(1.02,-.02),
                      yticklabels=['Blue','Purple','Red'] if i==0 else ['','',''], 
                      yticks=[1,0.5,0])
    plt.setp(ticks[1][1], rotation= 90, horizontalalignment='right',  verticalalignment='center')

    ax[1].set(ylim=(1.02,-.02), yticks=[])
    ax[2].set(ylim = mx_trait['growth_rate'])
    ax[3].set(ylim = mx_trait['growth_rate'])
    ax[4].set(ylim = mx_trait['interaction_inter'])
    ax[5].set(ylim = mx_trait['interaction_inter'])
    
    if i==0:
        ax[2].set_ylabel('Growth rate', )
        ax[0].set_ylabel('Collective colour', )
        ax[4].set_ylabel('Inter-colour competition', )
    else:
        ax[2].set_yticklabels([])
        ax[0].set_yticklabels([])
        ax[4].set_yticklabels([])
    titl = ax[1].set_title(title[i], )
    plt.setp(titl, position=(xcenter,plt.getp(titl, 'position')[1]))

    
    ###### ------------- ####
    ## Plot collective density 
    if True:
        cp_density = (data[i].data['cp_density'] * data[i].parameters['D'])
        # Before the break
        im = ax[0].contourf(cp_density[:,:Nbreak],
               extent=[0,Nbreak,0,1],
                levels=zbin_cp,
               colors=cmap_cp)
        # After the break. 
        im = ax[1].contourf(cp_density[:,Nbreak:],
                       extent=[Nbreak,cp_density.shape[1],0,1],
                        levels=zbin_cp,
                       colors=cmap_cp)
        # Color bar collectives
        if i == 0:
            cbar = fig.colorbar(im,ax=ax[0], cax=cax[0], ticks=tk)
            cbar.ax.set_ylabel('Number of collectives', )
            cbar.ax.set_yticklabels([x if x>0 else 0 for x in tk]) 

    ## Plot particle densities 
    CMAP_LIST = [plt.cm.Reds, plt.cm.Blues]
    if True:
        individual_traits_density = data[i].data['individual_traits_density']
        individual_traits_bins = data[i].data['individual_traits_bins']
        for pos,axid in zip((1,3), (2,4)):
            for color in (0,1):
                key = (ehs.COLOR_NAMES[color], ehs.POS_NAMES[pos])
                #cm = CMAP_LIST[color](np.arange(CMAP_LIST[color].N))
                #list_of_colors = CMAP_LIST[color](np.arange(CMAP_LIST[color].N))
                #list_of_colors[:,-1] = np.linspace(0,1,len( list_of_colors[:,-1]))
                list_of_colors = CMAP_LIST[color](np.logspace(0, CMAP_LIST[color].N, len(zbin)))
                list_of_colors[:,-1] = np.linspace(0,.8,len(list_of_colors[:,-1]))
    
                cm = ListedColormap(list_of_colors)
                    
                d = data[i].data['individual_traits'][key]
                d['c'] = d.value*d.number
                values = (d.groupby('generation').sum()['c']/d.groupby('generation').sum()['number']).values

                # Plot average 
                #ax[axid].plot(np.arange(0,Nbreak), 
                #              values[:Nbreak],
                #              color=['C3','C0'][color],
                #             ls='--', lw=1,)
                #ax[axid+1].plot(np.arange(Nbreak,individual_traits_density[key].shape[1]), 
                #              values[Nbreak:],
                #             color=['C3','C0'][color],
                #             ls='--',lw=1,)

                im = ax[axid].contourf(individual_traits_density[key][:,:Nbreak], 
                     extent=[0,
                             Nbreak,
                             individual_traits_bins[key[1]].min(),
                             individual_traits_bins[key[1]].max()], 
                             levels=zbin,
                                       antialiased=False,
                             colors=list_of_colors)
                im = ax[axid+1].contourf(individual_traits_density[key][:,Nbreak:], 
                                 extent=[Nbreak,
                                         individual_traits_density[key].shape[1],
                                         individual_traits_bins[key[1]].min(),
                                         individual_traits_bins[key[1]].max()], 
                                 levels=zbin,
                                         antialiased=False,
                                 colors=list_of_colors)

                if i == 0 and pos == 1:
                    cb = fig.colorbar(im,ax=ax[axid], cax=cax[1+color], ticks=zticks)
                    cax[1+color].set_ylabel('Number of {} individuals'.format(ehs.COLOR_NAMES[color])) 
                    cax[1+color].set_yticklabels(zbin_names) 

plt.tight_layout()
fig.savefig(os.path.join(FIGPATH,'full_trajectory.png'), bbox_inches='tight')
#fig.savefig(os.path.join(FIGPATH,'full_trajectory_hirez.png'), dpi=600, bbox_inches='tight')

# %% [markdown]
# Wide version (presentation)

# %% [code]
plt.rc('font', size=15)
fig = plt.figure(figsize=(16,9))
tk =  [1,100,1000]

#-------- Set the axis structure ---- ###
gs = [gridspec.GridSpec(nrows=3, ncols=5, left=0.05, right=0.48,
                        wspace=0.01, hspace=0.1),
      gridspec.GridSpec(nrows=3, ncols=5, left=0.51, right=0.96,
                         wspace=0.01, hspace=0.1)]
g = gridspec.GridSpec(nrows=3, ncols=1, left=0.97, right=0.99,
                 wspace=0.01, hspace=0.1)

cax = []
cax.append(fig.add_subplot(g[0, 0]))
cax.append(fig.add_subplot(g[1, 0]))
cax.append(fig.add_subplot(g[2, 0]))
# ---------#

xcenter = 0.41
title = ['Neutral Regime', 'Selective Regime']
for i in (0,1):

    ax = []
    ax.append(fig.add_subplot(gs[i][0, 0]))
    ax.append(fig.add_subplot(gs[i][0, 1:]))
    
    ax.append(fig.add_subplot(gs[i][1, 0]))
    ax.append(fig.add_subplot(gs[i][1, 1:]))
    
    ax.append(fig.add_subplot(gs[i][2, 0]))
    ax.append(fig.add_subplot(gs[i][2, 1:]))
    ylabel_font = fontdict={'fontweight': 'normal', 'size':'x-large'}

    for j,x in enumerate([1,3,5]):
        if j==2:
            lab = ax[x].set_xlabel('Collective generation',)
            plt.setp(lab, position=(xcenter,plt.getp(lab, 'position')[1]))
        ax[x].set(yticks=[],
                  xticks=[5000,10000],
                  xticklabels=[r'$5 \times 10^3$','$10^4$'] if j==2 else [])
        #ax[x].annotate(labels[i][j], **PANEL_DICT)
    for j,x in enumerate([0,2,4]):
        ax[x].set(xticks=[0,5,Nbreak],
                 xticklabels=[0,5,Nbreak] if j==2 else [])

    
    ticks = ax[0].set(ylim=(1.02,-.02), yticklabels=['Blue','Purple','Red'] if i==0 else ['','',''], yticks=[1,0.5,0])
    plt.setp(ticks[1][1], rotation= 90, horizontalalignment='right',  verticalalignment='center')

    ax[1].set(ylim=(1.02,-.02), yticks=[])
    ax[2].set(ylim = mx_trait['growth_rate'])
    ax[3].set(ylim = mx_trait['growth_rate'])
    ax[4].set(ylim = mx_trait['interaction_inter'])
    ax[5].set(ylim = mx_trait['interaction_inter'])
    
    if i==0:
        ax[2].set_ylabel('Growth rate', )
        ax[0].set_ylabel('Collective colour', )
        ax[4].set_ylabel('Inter-colour competition', )
    else:
        ax[2].set_yticklabels([])
        ax[0].set_yticklabels([])
        ax[4].set_yticklabels([])
    titl = ax[1].set_title(title[i], )
    plt.setp(titl, position=(xcenter,plt.getp(titl, 'position')[1]))

    
    ###### ------------- ####
    if True:
        cp_density = (data[i].data['cp_density'] * data[i].parameters['D'])
        individual_traits_density = data[i].data['individual_traits_density']
        individual_traits_bins = data[i].data['individual_traits_bins']

    ## Plot collective density 
    # Before the break
    
        im = ax[0].contourf(cp_density[:,:Nbreak],
               extent=[0,Nbreak,0,1],
                levels=zbin_cp,
               colors=cmap_cp)
        # After the break. 
        im = ax[1].contourf(cp_density[:,Nbreak:],
                       extent=[Nbreak,cp_density.shape[1],0,1],
                        levels=zbin_cp,
                       colors=cmap_cp)
        # Color bar collectives
        if i == 0:
            cbar = fig.colorbar(im,ax=ax[0],cax=cax[0], 
                         ticks=tk,
                         )
            cbar.ax.set_ylabel('Number of collectives', )
            cbar.ax.set_yticklabels([x if x>0 else 0 for x in tk]) 

    ## Plot particle densities 
    CMAP_LIST = [plt.cm.Reds, plt.cm.Blues]
    if True:
        for pos,axid in zip((1,3), (2,4)):
            for color in (0,1):
                key = (ehs.COLOR_NAMES[color], ehs.POS_NAMES[pos])
                cm = CMAP_LIST[color](np.arange(CMAP_LIST[color].N))

                d = data[i].data['individual_traits'][key]
                d['c'] = d.value*d.number
                values = (d.groupby('generation').sum()['c']/d.groupby('generation').sum()['number']).values

                cm = ListedColormap(cm)

                # Plot average 
                ax[axid].plot(np.arange(0,Nbreak), 
                              values[:Nbreak],
                              color=['C3','C0'][color],
                             ls='--', lw=1,)
                ax[axid+1].plot(np.arange(Nbreak,individual_traits_density[key].shape[1]), 
                              values[Nbreak:],
                             color=['C3','C0'][color],
                             ls='--',lw=1,)

                im = ax[axid].contourf(individual_traits_density[key][:,:Nbreak], 
                     extent=[0,
                             Nbreak,
                             individual_traits_bins[key[1]].min(),
                             individual_traits_bins[key[1]].max()], 
                             levels=zbin,
                             cmap=cm)
                im = ax[axid+1].contourf(individual_traits_density[key][:,Nbreak:], 
                                 extent=[Nbreak,
                                         individual_traits_density[key].shape[1],
                                         individual_traits_bins[key[1]].min(),
                                         individual_traits_bins[key[1]].max()], 
                                 levels=zbin,
                                 cmap=cm)

                if i == 0 and pos == 1:
                    cb = fig.colorbar(im,ax=ax[axid],
                                      cax=cax[1+color],
                                      ticks=zbin)
                    #cax[1+color].set_ylabel('Number of {} individuals'.format(ehs.COLOR_NAMES[color]),
                     #                       ) 
                    cax[1+color].set_yticklabels(zbin_names) 

plt.tight_layout()
#fig.savefig(os.path.join('.','full_trajectory.png'),dpi=600, bbox_inches='tight')
#fig.savefig(os.path.join(FIGPATH,'full_trajectory_hirez.png'), dpi=600, bbox_inches='tight')

# %% [code]


# %% [code]


# %% [code]


# %% [code]


# %% [code]


# %% [code]
````
