## Supplementary material for "Eco-evolutionary dynamics of nested Darwinian populations and the emergence of community-level heredity": Python code, figures and simulations: full_trajectory.py

```
#!/usr/bin/env python3
""" full_trajectory.py -- main entrypoint to start a simulation of the heredity model.
Copyright 2018 Guilhem Doulcier, Licence GNU GPL3+
"""

from functools import partial
import argparse
import multiprocessing
import datetime
import platform
import sys

import numpy

import estaudel.stochastic as stochastic
import estaudel.escaffolding as escaffolding
import estaudel.heredity.stochastic as model
import estaudel.heredity.process

# Mapping Arguments: (default_value, documentation)
PARAMETERS = {
    'D': (1000, 'Number of collectives'),
    'B': (15, 'Bottleneck size'),
    'N': (10000, 'Number of generations,'),
    'NPROC': (2, 'Number of CPU to use.'),
    'T': (1, 'Length of growth phase,'),
    'steps': (100, 'Number of discrete steps for bdm process.'),
    'skip': (100, ' Frequency at which growth trajectory are saved.'),
    'mutation_rate': (1, 'Probability that a birth event will also be a mutation event.'),
    'mutation_effect': ({1: 0.1, 3: 0.1}, 'Amplitude of mutational effects {trait:amplitude}'),
    'carrying_capacity': (1500, 'Max number of particle in a collective (will divide the a_intra, a_inter).'),
    'collectiveSelectionStrength': (1.0, ' Localisation parameter for the collective fitness function.'),
    'max_types': (4, 'Maximum number of types per collective'),
    'goal': (.5, ' Optimal proportion of types.'),
    'initial_type0': ((0, 6, .8, .15), 'intial traits (c, r, a_intra, a_inter)'),
    'initial_type1': ((1, 4, .3, .15), 'initial traits (c, r, a_intra, a_inter)'),
    'percentile': (20, ' Percentile of collective that go extinct each generation'),
    'selection': ('rank', 'Rank or neutral'),
    'continue': (None, 'Path to an output file from which extract the initial population'),
    'name': (None, 'Base name for output'),
    'force_traits': (False, 'When continue, Reset traits to initial_type.')
}


def empty_func(x):
    return None


def main(p, pool=None, filename=None):
    """ Assemble elements of the model and run the simulation. """
    ####  Mutation ####
    # Create a function that maps the phenotype before mutationm to the phenotype
    # after mutation. (phenotype->phenotype)
    if len(p['mutation_effect']):
        mutation_function = partial(stochastic.normal_mutation_abs,
                                    effect=p['mutation_effect'])
    else:
        print('No mutation !')
        mutation_function = empty_func
        p['mutation_rate'] = 0

    #### Growth dynamics ####
    # Create a function simulating the internal ecology of a collective.
    # dil_and_growth simulates the dilution of the parent, followed by the
    # growth of the new collective.
    growth = partial(stochastic.discrete_bdm_process,
                     T=p['T'],
                     steps=p['steps'],
                     skip=p['skip'],
                     mutation_rate=p['mutation_rate'],
                     mutation_function=mutation_function,
                     rate_function=partial(model.bd_rates, K=p['carrying_capacity']))

    dil_and_growth = partial(escaffolding.dilution_and_growth,
                             bottleneck_size=p['B'],
                             growth_function=growth)

    ### Collective Selection functions ###
    # collective_fitness_func associates to each collective a "quality"
    # collective_birth_death_func performs the selection.
    collective_fitness_func = partial(model.collective_fitness,
                                      var=p['collectiveSelectionStrength'],
                                      goal=p['goal'])

    if p['selection'] == 'neutral':
        collective_birth_death_func = escaffolding.collective_birth_death_neutral
    elif p['selection'] == 'rank':
        collective_birth_death_func = escaffolding.collective_birth_death_process
    elif p['selection'] == 'soft':
        collective_birth_death_func = escaffolding.collective_birth_death_process_soft
    elif p['selection'] == 'serial':
        collective_birth_death_func = escaffolding.collective_serial_transfer

    collective_birth_death_func = partial(collective_birth_death_func,
                                          percentile=p['percentile'])

    ### Prepare the initial conditions ####
    # Initial condition is pop, a list of (phenotype, state)
    # If continue is a filename, the simulation will use the last generation of this file
    # as initial condition.
    if p['continue'] is not None:
        out = escaffolding.Output(0, 0, 0, 0)
        out.load(p['continue'])
        N = out.current_gen
        print('Loading gen {} from {}'.format(N, p['continue']))
        if p['force_traits']:
            pop = [(model.gen_collective(p['max_types'], p['B'], p['initial_type0'], p['initial_type1'])[0],
                    out.state[:, N, d])
                   for d in range(out.state.shape[2])]
        else:
            pop = [(out.phenotype[:, :, N, d], out.state[:, N, d])
                   for d in range(out.state.shape[2])]
    else:
        pop = [model.gen_collective(p['max_types'], p['B'], p['initial_type0'], p['initial_type1'])
               for i in range(p['D'])]

    ### Prepare the output object ###
    # The output object store the trajectory of the model.
    output = escaffolding.Output(p['N'], p['D'], pop[0][0].shape[0], pop[0][0].shape[1])
    output.parameters = p

    ### Run the simulation ###
    output, pop = escaffolding.collective_generations(p['N'], pop, output,
                                                      collective_fitness_func,
                                                      collective_birth_death_func,
                                                      dil_and_growth, filename=filename, pool=pool)
    sys.stdout.flush()

    ### Extract some interesting statistics from the trajectory.###
    estaudel.heredity.process.extract(output, pool)
    sys.stdout.flush()

    # Save the output as a pkle file for further analysis.
    if filename is not None:
        output.save(filename+'.pkle')

    try:
        import pandas
        import numpy as np
        distrib_cp = output.data['cp_value'][output.current_gen, :]
        stats = {'mean_cp': distrib_cp.mean()}
        for k, v in output.data['individual_traits'].items():
            tr = v[v.generation == v.generation.max()]
            stats[k] = np.sum(tr.value * tr.number)/tr.number.sum()
        stats.update(output.parameters)
        pandas.DataFrame([stats]).to_csv(filename+'.csv')
    except Exception as ex:
        print(ex)
    return output


def cli_interface():
    """Command line interface for the heredity model"""

    # Prepare the argument parser with one argument per parameter.
    parser = argparse.ArgumentParser(description='Nested Darwinian Population simulation')
    for name, (_, man) in PARAMETERS.items():
        parser.add_argument('--'+name, help=man, nargs="?")
    args = parser.parse_args()

    # Update the parameters values from the arguments
    # UNSAFE USE OF EVAL ! Be Careful :-)
    param = (
        {k: eval(v) if v is not None else PARAMETERS[k][0] for k, v in vars(args).items()})

    # Start to keep track of time to compute the duration of the simulation.
    tstart = datetime.datetime.now()
    time_str = tstart.strftime('%Y%m%d_%H%M%S')

    # Setup the multiprocessing pool with the right number of parameters.
    # Intializiser np.random.seed ensures that the workers have a different RNG seed.
    pool = multiprocessing.Pool(param['NPROC'], initializer=numpy.random.seed)

    # Generate a filename
    filename = ((param['name'] + "_") if param['name'] is not None else '')
    filename += time_str + "_"
    filename += format_arguments(args)

    # Welcome message.
    print('='*80)
    print("{: ^80}".format("Stochastic Simulation of Nested Darwinian Populations"))
    print('='*80)
    print('Started at {}'.format(time_str))
    print('On {} | Python {}'.format(platform.platform(), platform.python_version()))
    print('Using {} / {} CPU'.format(param['NPROC'], multiprocessing.cpu_count()))
    print("{:=^80}".format(' Parameters '))
    print("\n".join(sorted(["{}: {}".format(k, v) for k, v in param.items()])))
    print("Will save in {}".format(filename))

    # Run the simulation
    main(param, pool, filename)

    # Close the multithreading pool
    pool.close()
    pool.join()

    # Display a last message with the time elapsed.
    print('='*80)
    tend = datetime.datetime.now()
    print('Ran in {}'.format((tend-tstart).total_seconds()))


def format_arguments(args):
    """Generate a filename-compatible string from the arguments"""
    def fmt(s):
        """ Replace a bunch of characters"""
        for c in '{}:"_()':
            s.replace(c, '')
        s.replace(',', '_')
        return s

    args_fmt = ["{}{}".format(k, fmt(v))
                for k, v in vars(args).items()
                if (k != 'name' and v is not None)]

    return "_".join(args_fmt)


if __name__ == "__main__":
    cli_interface()
```
