## Supplementary material for "Eco-evolutionary dynamics of nested Darwinian populations and the emergence of community-level heredity": Python code, figures and simulations: noselection_nomutation.ipynb

````
# %% [code]
import os 
import numpy as np
import matplotlib.pyplot as plt
import matplotlib.gridspec as gridspec

import estaudel.escaffolding as escaffolding
import estaudel.heredity.display 

from config import FIGPATH, PANEL_DICT, FONT_DICT, SUPFIGPATH

# %% [markdown]
Here are the commands to generate the data: 

Continuing the trajectory from `full.pkle` without mutation nor selection. 

```bash
./full_trajectory.py --N 1000 --continue \"full.pkle\" --NPROC 8 --selection \"neutral\" --name \"cont\" --mutation_rate 0
```

Same, but forcing a reset of the traits values: 

```bash
./full_trajectory.py --N 1000 --continue \"full.pkle\" --NPROC 8 --selection \"neutral\" --name \"cont\" --mutation_rate 0  --force_traits True
```

# %% [code]
path_derived = 'data/cont/cont_20190426_080331_NPROC8_mutation_rate0_N1000_continue"full.pkle"_selection"neutral".pkle'
path_intial = 'data/cont/cont_20190426_084814_mutation_rate0_selection"neutral"_N1000_force_traitsTrue_NPROC8_continue"full.pkle".pkle'

# %% [code]
data_derived = escaffolding.load(path_derived)
data_initial = escaffolding.load(path_intial)
for d in [data_initial, data_derived]:
    print(d)
    print("\n".join(["{}: {}".format(k,v) for k,v in sorted(d.parameters.items())]))
    print('-'*80)

# %% [code]
fig = plt.figure(figsize=(7,4))

# Axes grid spec. 
grid_right = gridspec.GridSpec(nrows=1, ncols=5, left=0.0, right=0.96, wspace=0.01, hspace=0.3)
grid_colorbar = gridspec.GridSpec(nrows=1, ncols=1, left=0.97, right=0.99, wspace=0.01, hspace=0.3)

# Create the axes
cax = fig.add_subplot(grid_colorbar[0, 0])
#ax_left = fig.add_subplot(grid_left[0, 1:])
#ax_left_zoom = fig.add_subplot(grid_left[0, 0])
ax_right = fig.add_subplot(grid_right[0, 1:])
ax_right_zoom = fig.add_subplot(grid_right[0, 0])

# Label the panels 
#ax_right.annotate('B', **PANEL_DICT)
#sup = fig.suptitle('Without collective selection nor particle mutation', fontsize='x-large')

estaudel.heredity.display.plot_density(data_derived, ax_right, zoom_first=10, 
             title='Derived particle traits',    colorbar=True, 
             cax=cax,
             ax_zoom=ax_right_zoom)

#plt.savefig(os.path.join(FIGPATH, 'nomutation_norselection2.png'), dpi=900, bbox_inches='tight')

# %% [code]
plt.rc('font', size=15)
fig = plt.figure(figsize=(15,4))

# Axes grid spec. 
grid_left =  gridspec.GridSpec(nrows=1, ncols=5, left=0.05, right=0.48, wspace=0.01, hspace=0.3)
grid_right = gridspec.GridSpec(nrows=1, ncols=5, left=0.51, right=0.96, wspace=0.01, hspace=0.3)
grid_colorbar = gridspec.GridSpec(nrows=1, ncols=1, left=0.97, right=0.99, wspace=0.01, hspace=0.3)

# Create the axes
cax = fig.add_subplot(grid_colorbar[0, 0])
ax_left = fig.add_subplot(grid_left[0, 1:])
ax_left_zoom = fig.add_subplot(grid_left[0, 0])
ax_right = fig.add_subplot(grid_right[0, 1:])
ax_right_zoom = fig.add_subplot(grid_right[0, 0])

# Label the panels 
PANEL_DICT.update({'xy':(0.95,0.1), 'size':20})
ax_left.annotate('A', **PANEL_DICT)
ax_right.annotate('B', **PANEL_DICT)
#sup = fig.suptitle('Without collective selection nor particle mutation', fontsize='x-large')

# Plot density
estaudel.heredity.display.plot_density(data_initial, ax_left, zoom_first=10,
             title='Initial particle traits', 
             ax_zoom=ax_left_zoom,
             colorbar=True, 
             cax=cax)
estaudel.heredity.display.plot_density(data_derived, ax_right, zoom_first=10, 
             title='Derived particle traits', 
             ax_zoom=ax_right_zoom)
ax_right_zoom.set(yticklabels=[], ylabel='')
plt.savefig(os.path.join(FIGPATH, 'nomutation_norselection.png'), bbox_inches='tight')

# %% [markdown]
## noselection but mutations

# %% [code]
path = 'cont_20190425_084340_continue"full.pkle"_NPROC8_N1000_selection"neutral".pkle'
data = escaffolding.load(path)

# %% [code]
fig, ax = plt.subplots(1,1,figsize=(10,4))
estaudel.heredity.display.plot_density(data, ax=ax, colorbar=True)
ax.set_xlim(0,300)
ax.set_title('Derived traits, without collective selection, with particle mutations.')
plt.savefig(os.path.join(SUPFIGPATH,'noselection.png'))

# %% [markdown]
# Selection but no mutations
This is the textbook "Stochastic corrector" mechanism

```
python3 full_trajectory.py --mutation_rate 0 --mutation_effect "{}" --N 1000 --D 1000 --initial_type1 "(1,5,0.7,0.3)" 
```

# %% [code]
path = "20200409_153204_D1000_N1000_mutation_rate0_mutation_effect{}_initial_type1(1,5,0.7,0.3).pkle"
data = escaffolding.load(path)

plt.rc('font', size=14)
fig = plt.figure(figsize=(19,8))
fig, ax = plt.subplots(1,1,figsize=(10,4))
estaudel.heredity.display.plot_density(data, ax=ax, colorbar=True)
ax.set_xlim(0,1000)
ax.set_title('Ancestral traits,\n with collective selection, without particle mutations.')
plt.savefig(os.path.join(SUPFIGPATH,'nomut_with_selection.png'), bbox_inches='tight')
````
