## Supplementary material for "Eco-evolutionary dynamics of nested Darwinian populations and the emergence of community-level heredity": Python code, figures and simulations: particle_ecology.ipynb

```
# %% [markdown]
# Particle ecology

# %% [code]
from functools import partial
import numpy as np
import matplotlib.pyplot as plt
from mpl_toolkits.axes_grid1 import make_axes_locatable
import scipy.integrate

import estaudel.escaffolding as escaffolding
import estaudel.stochastic as stochastic
import estaudel.heredity.process as ehp
import estaudel.heredity.deterministic as ehd
import estaudel.heredity.stochastic as ehs
from config import FIGPATH, PANEL_DICT, FONT_DICT, SUPFIGPATH
import os

# %% [markdown]
# Data
Here we extract the data from the full simulation in order to get the right traits values.

# %% [code]
generations_to_extract = [3,9000]
tstep = 500
path = "data/full_trajectory/full_trajectory.321.1_20180629_163801_selection\"rank\"_NPROC8.pkle"

# %% [code]
out = escaffolding.load(path)
out.parameters

# %% [markdown]
# Doing small simulations

In general I do not record the full trajectories of all the collectives (it would take too much space) so this figure is made by doing a new simulation, using the resident traits and initial conditions that where extracted from the full simulation.

We turn off the mutations in this simulation to not overload the figure.

# %% [code]
np.random.seed(120292)
traj = {}
traj_sto = {}
lv_param = {}
time = np.linspace(0,out.parameters['T'],tstep)
N_simulations = 300
for n in generations_to_extract:
    traj[n] = []
    traj_sto[n] = []

    for x,P in [(out.data['cp_value'][n-1,out.parents[n-1][i]], 
                 np.array(out.data['resident_pheno'][out.data['resident_id'][n,i]]))
                for i in range(N_simulations)]:           
        # Initial conditions
        state = np.array(np.array([out.parameters['B']*(1-x),out.parameters['B']*x]))
        
        # Deterministic trajectory 
        r,a = ehd.convert_phenotypes_to_lv(P, out.parameters['carrying_capacity'])
        func = partial(ehd.lotka_volterra,r=r,a=a)
        traj[n].append(scipy.integrate.odeint(func,
                                y0=state,
                                t=time))
        lv_param[n] = (r,a)
        # Stochastic trajectory
        rates = partial(ehs.bd_rates,K=out.parameters['carrying_capacity'])
        growth_no_mutation = partial(
            stochastic.discrete_bdm_process,
            T=out.parameters['T'],
            steps=tstep,
            skip=1,
            mutation_rate=0,
            mutation_function=lambda x:None,
            rate_function=rates)
        traj_sto[n].append(growth_no_mutation(state=state, phenotypes=P)['trajectory'].transpose())

# %% [markdown]
# Plot 

# %% [code]
# select the highlighted collective
highlighted = {}

# For the first one we want something that overshoot at the end. 
phen = [t[-1,0] /t[-1,:].sum() for t in traj_sto[generations_to_extract[0]]]
highlighted[generations_to_extract[0]] = [(n,x) for n,x in sorted(enumerate(phen), key=lambda k:k[1])][3][0]

# For the last one we want something that overshoot in the middle
phen = [t[tstep//3,0] /t[tstep//3,:].sum() for t in traj_sto[generations_to_extract[1]]]
highlighted[generations_to_extract[1]] = [(n,x) for n,x in sorted(enumerate(phen),key=lambda k:k[1])][1][0]
highlighted

# %% [markdown]
## Particle Trajectories
This figure was made to give a first visual overview of the intra-collective particle ecology. We show the diversity of collective trajectories at the begining and end of the evolutionary trajectory. Then in a second panel, we zoom out on a single panel to show the trajectories of each type of particles. It is a good time to show how the ODE approximate the stochastic model. 

# %% [code]
s = out.parameters['collectiveSelectionStrength']
ftnes = lambda x :  1/np.sqrt(np.pi*2*s) * np.exp(-(0.5-x)**2 / 2*s)
ftnes_inv = lambda f: np.roots([1,
                                -1,
                                0.25+2*s*np.log(f*np.sqrt(2*np.pi*s))])
np.testing.assert_almost_equal(ftnes_inv(ftnes(0.77)).max(), 0.77)

# %% [code]
from mpl_toolkits.axes_grid1.inset_locator import inset_axes
from sklearn.neighbors import KernelDensity
import matplotlib.path as mpath
import matplotlib.patches as mpatches
import matplotlib.pyplot as plt
from matplotlib.patches import FancyArrowPatch, PathPatch
import matplotlib.gridspec as gridspec

# %% [code]
def plot_individual_dynamics(gen, ax_color, ax_highlight, ax_distrib, threshold=False):
    i = 3
    d = highlighted[gen]
    ### First panel: all collectives ###
    trmax = 250
    for sto in traj_sto[gen][:trmax]:
        ax_color.plot(time, sto[:,0]/sto.sum(1), color=(.8,.8,.8),lw=.3)
    ax_color.plot(time, traj_sto[gen][d][:,0]/traj_sto[gen][d].sum(1), color='C1')
    
    phen = [t[-1,0] /t[-1,:].sum() for t in traj_sto[gen]]
    f = [ftnes(x) for x in phen]
    
    if threshold:
        threshold = np.percentile(f, out.parameters['percentile'])
        ax_color.fill_between([0.97,1], [ftnes_inv(threshold)[0]]*2,[ftnes_inv(threshold)[1]]*2, 
                             color='C2', zorder=99, alpha=.2)
        ax_color.hlines(ftnes_inv(threshold),0.97,1, color='C2', zorder=99)
    ax_color.set(xlim=(0,1), ylim=(-0.02,1.02),title='Generation {}'.format(gen))

    inset_distrib(phen, ax_distrib, out.parameters['percentile'])
    
    ### Second Panel: zooming on a single collective ###
    # Stochastic trajectory with a full line
    ax_highlight.semilogy()
    ax_highlight.plot(time, traj_sto[gen][d][:,0], color='C3')
    ax_highlight.plot(time, traj_sto[gen][d][:,1], color='C0')
    
    # ODEs with a dotted line
    ax_highlight.plot(time, traj[gen][d][:,0], color='C3',ls=':')
    ax_highlight.plot(time, traj[gen][d][:,1], color='C0',ls=':')
    
    # Legends and axis labels
    ax_highlight.vlines(0,0,0, linestyles=':', label='ODE approximation')
    ax_highlight.vlines(0,0,0, linestyles='-', label='Stochastic process')
    ax_highlight.legend()
    
    ax_highlight.set(xlabel='Time', xlim=(0,1), ylim=(1,4500))

    
def inset_distrib(data, ax, percentile, exact_percentile=False):
    kde = KernelDensity(kernel='gaussian', bandwidth=0.02).fit(np.array(data).reshape(-1, 1))
    ax.set(xticks = [], yticks=[0,0.5,1], ylim=(0,1), yticklabels=[])
    # Dark - distrib
    x = np.linspace(0,1,1000)[:, np.newaxis]
    log_dens = kde.score_samples(x)
    ax.fill(np.concatenate([[0], np.exp(log_dens).flatten(),[0]]),
            np.concatenate([[0],x.flatten(),[1]]), 
            alpha=1, color='grey')
    f = [ftnes(x) for x in data]
    
    # Green - selected part
    threshold = np.percentile(f, percentile)
    low,high = ftnes_inv(threshold)
    
    # Ensure that the grey area is still visible even if its very small.
    # We are aiming for clarity not corectness here
    if not exact_percentile:
        low = max(0.05, min(0.96, low))
        high = min(0.96, max(0.05,high))
        print(low,high)

    x = np.linspace(low,high,1000)[:, np.newaxis]
    log_dens = kde.score_samples(x)
    ax.fill(np.concatenate([[0], np.exp(log_dens).flatten(),[0]]), 
            np.concatenate([[low],x.flatten(),[high]]),
            color='C2')

# %% [code]
plt.rc('font', size=15)
fig = plt.figure(figsize=(13,6))

# Axes grid spec. 
grid_left =  gridspec.GridSpec(nrows=2, ncols=15, left=0.05, right=0.48, wspace=0.01, hspace=0.1)
grid_right = gridspec.GridSpec(nrows=2, ncols=15, left=0.51, right=0.96, wspace=0.01, hspace=0.1)

ax_color1 = fig.add_subplot(grid_left[0, :-1])
ax_zoom1 = fig.add_subplot(grid_left[0, -1])
ax_highlight1 = fig.add_subplot(grid_left[1, :-1])


ax_color2 = fig.add_subplot(grid_right[0, :-1])
ax_zoom2 = fig.add_subplot(grid_right[0, -1])
ax_highlight2 = fig.add_subplot(grid_right[1, :-1])

plot_individual_dynamics(generations_to_extract[0], ax_color1, ax_highlight1, ax_zoom1)
plot_individual_dynamics(generations_to_extract[1], ax_color2, ax_highlight2, ax_zoom2)

ax_highlight1.set(ylabel='Number of\n Individuals')        
ax_color1.set(ylabel='Proportion of\n {} particles'.format(ehs.COLOR_NAMES[0]))
ax_color1.set(ylim=(0,1),yticks=[0,0.5,1], yticklabels=[0,0.5,1])
ax_color2.set(ylim=(0,1), yticks=[0,0.5,1], yticklabels=[]);

pd = PANEL_DICT
if 'xy' in PANEL_DICT:
    del pd['xy']
ax_color1.annotate('A',xy=(0.05, 0.1),**pd)
ax_color2.annotate('B',xy=(0.05, 0.1),**pd)
ax_highlight1.annotate('C',xy=(0.05, 0.1),**pd)
ax_highlight2.annotate('D',xy=(0.05, 0.1),**pd)


for a in (ax_color1, ax_highlight1, ax_color2, ax_highlight1):
    a.set(xticks=[0,0.2,0.4,.6,.8,1], xticklabels=[0,.2,.4,.6,.8,1])
ax_highlight2.set(yticklabels=[])
ax_color2.set(yticklabels=[])
ax_color1.set(xticklabels=[])
ax_color2.set(xticklabels=[])

ax2 = ax_zoom2.twinx()
ax2.set_ylim(0,1)
ticks = ax2.set(ylim=(-0.02,1.02), yticklabels=['Blue','Purple','Red'], yticks=[0,0.5,1])
plt.setp(ticks[1][1], rotation= -90, horizontalalignment='left',  verticalalignment='center')
ax2.plot([],[]);

plt.tight_layout()
fig.savefig(os.path.join(FIGPATH,'particle_ecology.pdf'), bbox_to_inches='tight')

# %% [markdown]
## Empirical G's

# %% [code]
from config import BLUE_RED

# %% [code]
np.random.seed(120292)
empirical_g = {}
time = np.linspace(0,out.parameters['T'],tstep)
N_simulations = 10
replicates = 50
xlist = np.arange(out.parameters['B']+1)

rates = partial(ehs.bd_rates,K=out.parameters['carrying_capacity'])

growth_no_mutation = partial(
            stochastic.discrete_bdm_process,
            T=out.parameters['T'],
            steps=tstep,
            skip=1,
            mutation_rate=0,
            mutation_function=lambda x:None,
            rate_function=rates)

for n in generations_to_extract:
    empirical_g[n] = {}
    P = np.array(out.data['resident_pheno'][out.data['resident_id'][n,highlighted[n]]])
    for x in xlist:
        prop = x/out.parameters['B']
        empirical_g[n][prop] = []
        for _ in range(replicates):
            # Initial conditions
            state = np.array(np.array([x,out.parameters['B']-x]))
            endstate = growth_no_mutation(state=state, phenotypes=P)['trajectory'][:,-1]
            empirical_g[n][prop].append(endstate[0]/endstate.sum())

# %% [code]
gfunc = {}
gg = {}
plist = np.linspace(0,1,500)
fixed_points = {}
deter_traj = {}
for n in generations_to_extract:
    P = np.array(out.data['resident_pheno'][out.data['resident_id'][n,highlighted[n]]])
    r,a = ehd.convert_phenotypes_to_lv(P, out.parameters['carrying_capacity'])
    g = ehd.get_gfunc(r=r,A=a, B=out.parameters['B'], T=out.parameters['T'])
    gfunc[n] =  [g(p) for p in plist]

    deter_traj[n], gg[n] = ehd.iterate_ecology(r, a, B=out.parameters['B'], T=out.parameters['T'], x0=1/out.parameters['B'])
    if n == 9000:
        fixed_points[n] = [(0, False), (scipy.optimize.fsolve(lambda x: g(x)-x, 0.5), True), (1,False)] 
    else:
        fixed_points[n] = [(0, False),(1,True)]

# %% [code]
plt.rc('font', size=15)
fig, ax = plt.subplots(1,2,figsize=(7.5,4))
for i, gen in enumerate(generations_to_extract):
    for x,values in empirical_g[gen].items():
        ax[i].plot(plist,  gfunc[gen], color='C1', zorder=0)
        
        ax[i].scatter(x, np.mean(values), marker='o', color=BLUE_RED(np.mean(values)), zorder=2)
        ax[i].errorbar(x, np.mean(values), np.std(values),  color=BLUE_RED(np.mean(values)),zorder=2)
        ax[i].set(xlim=(-0.01,1.01),ylim=(-0.01,1.01))
        ax[i].set(title='Generation {}'.format(gen))

    for fp, stability in fixed_points[gen]:
        ax[i].scatter(fp,fp, color='C1',# if stability else 'w',
                     # if stability else 'C1',
                     marker='x', s=100, zorder=1)

    ax[i].vlines(gg[gen][0],0,gg[gen][1],color='k', alpha=0.5, linestyles='-.')
    for n,(p,p1) in enumerate(zip(gg[gen][:-1],gg[gen][1:])):
        #ax[i].vlines(p,0,p1,color='C4', alpha=0.4)
        ax[i].hlines(p1,p,p1,color='k', linestyles='-.',alpha=.5)
        if n+2<len(gg[gen]):
            ax[i].vlines(p1,gg[gen][n+2],p1,color='k',alpha=.5, linestyles='-.')

    ax[i].plot([0,1],[0,1], color='k', ls='-', alpha=.1)

bbox_legend2 = dict(xy=(.08, .92), xycoords="axes fraction",
               ha='center', va='center',
                bbox=dict(boxstyle="round",
                   ec=(.1, 0.1, .1),
                   fc=(1., 1, 1),
               ))
for i in (0,1):
    ax[i].set(xticks=[0,.25,.5,.75,1], xticklabels=[0,'',0.5,'',1])
    ax[i].set(yticks=[0,.25,.5,.75,1], yticklabels=[0,'',0.5,'',1])
    ax[i].set(xlabel='Newborn colour')
ax[0].set(ylabel='Adult colour')
ax[1].set(yticklabels=['','','','',''])
ax[0].annotate('A',**bbox_legend2)
ax[1].annotate('B',**bbox_legend2)
plt.tight_layout()
fig.savefig(os.path.join(FIGPATH,"empirical_gfunc.pdf"), bbox_inches="tight")

# %% [markdown]
## G function trajectory

# %% [code]
# Rewind the ancestry of a collective
id0 = 1
idx = [1]
for par in out.parents[::-1]:
    idx.append(par[idx[-1]])
idx = idx[::-1] # now they are in increasing order

# %% [code]
all_g = {}
plist = np.linspace(0,1,500)
gtraj_gen = np.arange(0,9001,25)
for gen in gtraj_gen:
    col = idx[gen]
    P = np.array(out.data['resident_pheno'][out.data['resident_id'][gen,col]])
    r,a = ehd.convert_phenotypes_to_lv(P, out.parameters['carrying_capacity'])
    g = ehd.get_gfunc(r=r,A=a, B=out.parameters['B'], T=out.parameters['T'])
    all_g[gen] = [g(p) for p in plist]

# %% [code]
L = 8
sp = 1/20
plt.rc('font', size=10)
fig, axes = plt.subplots(L,1,figsize=(12,.75*L))
per_line = 1+np.max(gtraj_gen)//L
tick = {j:[] for j in range(L)}
ticklabel = {j:[] for j in range(L)}
scale = 0.80
for i in gtraj_gen:
    j = i//per_line
    if j>=8:
        print(i,j)
    x_shift = j*per_line+(i%per_line)*sp
    axes[j].plot(scale*plist+x_shift, scale*plist, color='k', alpha=.2)
    axes[j].plot(scale*plist+x_shift, scale*np.array(all_g[i]), color='C0')
    if i%250==0:
        tick[j].append(scale*0.5+x_shift)
        ticklabel[j].append(i)
        
for j,ax in enumerate(axes):
    ax.spines['right'].set_color('none')
    ax.spines['top'].set_color('none')
    ax.spines['left'].set_color('none')
    ax.set(xlim=(j*per_line-0.5*scale,j*per_line+1+per_line*sp),
           ylim=(0,scale))
    ax.set(yticks=[],
          xticks=tick[j], xticklabels=ticklabel[j])
plt.tight_layout()
axes[-1].set_xlabel('Collective generation', fontdict={'size':16})
fig.savefig(os.path.join(FIGPATH,'trajectory_of_g.pdf'), bbox_inches='tight')

# %% [markdown]
# Suplementary : Iterations

# %% [code]
def plot_g(plist, gfunc, ax, gg):
    ax.plot(plist,  gfunc, color='C1', zorder=0)
    ax.set(xticks=[0,.5,1],yticks=[0,.5,1], xlabel='Initial colour', ylabel='Final colour')
    ax.plot([0,1],[0,1], color='k', alpha=0.1)
    ax.grid()
    ax.vlines(gg[0],0,gg[1],color='C2', alpha=0.8, linestyles='-.')

    for n,(p,p1) in enumerate(zip(gg[:-1],gg[1:])):
        ax.hlines(p1,p,p1,color='C2', linestyles='-.',alpha=.8)
        if n+2<len(gg):
            ax.vlines(p1,gg[n+2],p1,color='C2',alpha=.8, linestyles='-.')

    
def plot_vector_field(r, A, color='grey', text=True, ax=None, mx=None, B=None):
    """Plot the LV-vector field with null isoclines and equilibrium."""
    if ax is None:
        ax = plt.gca()
        
    # Flow 
    if mx is None:
        mx = np.max(1/A)
    dx = mx/30
    x = np.linspace(0,mx+dx,20)
    X,Y = np.meshgrid(x,x)
    
    flow = np.vectorize(lambda x,y: ehd.lotka_volterra([x,y],r=r,a=A),
                        signature='(),()->(2)')
    U = flow(X,Y)
    ax.streamplot(X, Y, U[:,:,0], U[:,:,1], density=1, color=color,linewidth=0.8)
    

    ax.set(xlabel='$N_0$',ylabel='$N_1$',
           xticks=[],yticks=[],
           xlim=(-dx,mx+dx), ylim=(-dx,mx+dx))

# %% [code]
plt.rc('font', size=15)
fig = plt.figure(figsize=(7.5,5))
labels = ['AB','CD']
for i, gen in enumerate(generations_to_extract):
    ax_phase = plt.subplot2grid((2,3),(i,2))

    #ax_g = plt.subplot2grid((2,4),(i,3))
    ax_traj = plt.subplot2grid((2,3),(i,0),1,2)
    #plot_g(plist, gfunc[gen], ax_g, gg[gen])
    nmax = np.concatenate(deter_traj[gen]).max()
    ax_phase.annotate(labels[i][1],**PANEL_DICT)
    ax_traj.annotate(labels[i][0],**PANEL_DICT)
    
    T = deter_traj[gen][0].shape[0]
    ax_traj.set(xlabel='Time',ylabel='N',
                xticks=[T*i for i in range(len(deter_traj[gen])+1)],
               xticklabels=[i for i in range(len(deter_traj[gen])+1)])
    #ax_traj.grid()
    for i,tj in enumerate(deter_traj[gen]):
        ax_traj.plot(np.arange(T)+i*T, tj[:,0], color='C3')
        ax_traj.plot(np.arange(T)+i*T, tj[:,1], color='C0')
        ax_phase.plot(tj[:,0],tj[:,1], color='C2')
    for i,(tj,tjp1) in enumerate(zip(deter_traj[gen][:-1],deter_traj[gen][1:])):
        ax_phase.plot([tj[-1,0],tjp1[0,0]],[tj[-1,1],tjp1[0,1]], ls=':', color='C2')
        ax_traj.plot([T*(i+1),T*(i+1)], [tj[-1,0],tjp1[0,0]], color='C3', ls='-.')
        ax_traj.plot([T*(i+1),T*(i+1)], [tj[-1,1],tjp1[0,1]], color='C0', ls='-.')

    plot_vector_field(lv_param[gen][0],lv_param[gen][1], mx=nmax, ax=ax_phase, color=(0,0,0,0.1))

fig.tight_layout()
fig.savefig(os.path.join(SUPFIGPATH,'deterministic_iterations.pdf'), bbox_inches='tight')
fig.savefig(os.path.join(SUPFIGPATH,'deterministic_iterations.png'), bbox_inches='tight')
```
