## Supplementary material for "Eco-evolutionary dynamics of nested Darwinian populations and the emergence of community-level heredity": Python code, figures and simulations: positive_interactions.ipynb

```
# %% [markdown]
# Positive interactions
Those figures extend the analysis of the deterministic model to mutualistic and predator-prey interactions. 

Mutualistic interaction requires particular care because since the pop size can grow unbounded, the integration quickly overflow. 

# %% [code]
from collections import OrderedDict, Counter
from multiprocessing import Pool
from functools import partial
import pickle

import numpy as np

import scipy.optimize
import scipy.integrate
import scipy.misc

# Matplotlib related imports 
import matplotlib.pyplot as plt
from matplotlib import rc
from matplotlib.colors import LinearSegmentedColormap
from matplotlib.colors import ListedColormap
from mpl_toolkits.axes_grid1.axes_divider import make_axes_locatable
from mpl_toolkits.axes_grid1.colorbar import colorbar
from mpl_toolkits.axes_grid1.inset_locator import inset_axes, zoomed_inset_axes
from mpl_toolkits.axes_grid1.anchored_artists import AnchoredSizeBar
from mpl_toolkits.axes_grid1.inset_locator import inset_axes
%matplotlib inline

import estaudel.escaffolding as escaffolding
import estaudel.heredity.process
import estaudel.heredity.deterministic as ed
import estaudel.heredity.deterministic as ehd

from estaudel.heredity.stochastic import COLOR_NAMES
from config import BLUE_RED, PANEL_DICT

# %% [code]
def pstar_background(ax, a0,a1,bar=True):
    plt.rc('font', size=17)
    xx = np.linspace(-1,1,200)
    Xa,Ya = np.meshgrid(xx,xx)

    f = np.vectorize(lambda x,y: ed.pstar(np.array([[a0,y], [x,a1]])))
    Za = f(Xa,Ya)

    ticks = [0,0.25,0.5,0.75,1]
    levels = np.linspace(0,1,20)

        
    # Background 
    im = ax.contourf(Xa, Ya, Za, cmap=BLUE_RED, levels=levels)
    ax.fill_between(x=[a0,1],y1=[a1,a1], y2=[1,1], color='#bab0ac')

    # Colorbar
    if bar:        
        div = make_axes_locatable(ax)
        cax = div.append_axes("right", size="4%", pad=0.11)
        cax.plot([0,1],[0.5,0.5],color='w',ls=':')
        cax.plot([0,1],[1,1],color='w',ls='-')
        cax.plot([0,1],[0,0],color='w',ls='-')
        cb = plt.colorbar(im, cax=cax, ticks=ticks)

    ax.hlines([0],-1,1,color='w',linestyles='-')
    ax.vlines([0],-1,1,color='w',linestyles='-')
    x = np.linspace(-1, a0)
    ax.plot(x, -(a0-a1)+x, color='w', ls=':')
    
def pstar_pred_prey(ax, a0 = 0.8, a1 = 0.4, bar=True):
    pstar_background(ax, a0=a0, a1=a1, bar=bar)
    # Opaque over non predation.
    ax.fill_between(x=[0,1],y1=[0,0], y2=[1,1], color='k', alpha=0.5)
    ax.fill_between(x=[-1,0],y1=[-1,-1], y2=[1,1], color='k', alpha=0.5)
    # Set axis
    ax.set(
            xlabel=r'$a_{10}$', 
            ylabel=r'$a_{01}$',
            #title=r'Stable fixed point of $G_\theta$ when $T \to \infty$',
            xlim=(-1,1),
            ylim=(-1,1),
            xticks=[0,a0-a1,a0,1],
            xticklabels=[0,'$a_{00}-a_{11}$','$a_{00}$',1,],
            yticklabels=[-1,0, '$a_{11}-a_{00}$','$a_{11}$',1,],
            yticks=[-1,0,a1-a0,a1,1]
    );

fig, ax = plt.subplots(1,2,figsize=(15,7))
pstar_pred_prey(ax[0],0.4,0.8,False)
ax[0].set_title('$a_{00}<a_{11}$')
ax[1].set_title('$a_{11}<a_{00}$')
pstar_pred_prey(ax[1],0.8,0.4,True)
plt.tight_layout()
plt.savefig('reviews/predator_prey.pdf', bbox_inches='tight')

# %% [code]
fig, ax = plt.subplots(1,1, figsize=(8,7))
a0 = 0.4
a1 = 0.8
pstar_background(ax, a0=a0, a1=a1)

ax.text(-1, -0.01, 'Mutualism', color='w', horizontalalignment='left', verticalalignment='top')
ax.text(-0.75, -0.75, 'Unbounded\n growth', color='w', horizontalalignment='center',
        fontdict={'style':'oblique', 'size':'small'},
        verticalalignment='center')
ax.text(1, 0.01, 'Competition', color='w', horizontalalignment='right', verticalalignment='bottom')
ax.text(1, -0.03, 'Predation\n (Red on Blue)', color='w', horizontalalignment='right', verticalalignment='top')
ax.text(-1, 0.03, 'Predation\n (Blue on Red)', color='w', horizontalalignment='left', verticalalignment='bottom')
ax.set(
        xlabel=r'$a_{10}$', 
        ylabel=r'$a_{01}$',
        title=r'Stable fixed point of $G_\theta$ when $T \to \infty$',
        xlim=(-1,1),
        ylim=(-1,1),
        xticks=[0,a0-a1,a0,1],
        xticklabels=[0,'$a_{00}-a_{11}$','$a_{00}$',1,],
        yticklabels=[-1,0, '$a_{11}-a_{00}$','$a_{11}$',1,],
        yticks=[-1,0,a1-a0,a1,1]
);


plt.tight_layout()
x = np.linspace(-(a0*a1),-1)
ax.plot(x,(a0*a1)/x, color='w')
plt.savefig('reviews/position_of_pstar_negative.pdf', bbox_inches='tight')

# %% [code]
def minipstar(ax, a0, a1):
    """Plot the colour at equilibrium as a function of a01, a10. 
    
        Args: 
            a0 (float): carrying capacity of type 0 individuals
            a1 (float): carrying capacity of type 1 individuals.
            ax (matplotlib.Axis): axis to plot on. 
        Returns:
            ax 
    """
    x = np.linspace(-1,a0, 10)
    ax.plot(x, a1-a0+x, color='white')
    ax.fill_between(x=[a0,1],y1=[a1,a1], y2=[1,1], color='#bab0ac')
    ax.fill_between(x=[a0,1],y1=[-1,-1], y2=[a1,a1], color=BLUE_RED(1.0))
    ax.fill_between(x=[-1,a0],y1=[a1,a1], y2=[1,1], color=BLUE_RED(0.0))
    ax.set(xticks=[0,a0,1],
            xticklabels=[0,'$a_{00}$',1,],
            yticklabels=[-1, 0, '$a_{11}$',1,],
            yticks=[-1,0,a1,1],
            xlabel='$a_{10}$',
            ylabel='$a_{01}$', 
            xlim=(-1,1), 
            ylim=(-1,1))
    Xa,Ya = np.meshgrid(np.linspace(-1,a0-0.001,10),
                        np.linspace(-1,a1-0.001,10))
    f = np.vectorize(lambda x,y: ed.pstar(np.array([[a0,y],[x,a1]])))
    Za = f(Xa,Ya)
    levels = np.linspace(0,1,20)
    im = ax.contourf(Xa, Ya, Za, cmap=BLUE_RED, levels=levels)
    x = np.linspace(0,-1)[1:]
    ax.plot(x,(a0*a1)/x, color='w')
    return ax

# %% [code]
def plt_traj(a0,a1, a01, a10, inset=None, ax=None, B=0.001):
    if ax is None:
        ax = plt.gca()
    A = np.array([[a0,a01],
                  [a10,a1]])
    r = np.array([40,40])
    tspan = np.linspace(0,1 if a0*a1>a01*a10 else 0.2, 1000)
    flow = partial(ed.lotka_volterra, r=r, a=A)
    traj = scipy.integrate.odeint( flow,[10,10], tspan )
    ax.plot(tspan, traj[:,0], color='C3')
    ax.plot(tspan, traj[:,1], color='C0')
    ax.hlines([1/a0, 1/a1], 0,tspan.max(),
              colors=['C3','C0'], linestyle='--')
    
    if inset:
        axin = inset_axes(ax, width=0.9, height=0.9, loc='center right')
        minipstar(axin, a0,a1)
        axin.scatter(a10,a01, color='w',s=3,marker='X')
    ax.set(xlabel='Time', xticks=[], title=str(A)+' \n x='+str(traj[-1,:]), yticks=[1/a0,B,1/a1],
                yticklabels=['$a_{00}^{-1}$','B','$a_{11}^{-1}$'])

# %% [code]
fig, ax = plt.subplots(2, 6, figsize=(20,6.66))
a = [(0.5,0.8), 
     (0.8,0.5)]
r = [30,20]
B = 0.0001
tspan = np.linspace(0,1,10000)
for i, (a0,a1) in enumerate(a):
    minipstar(ax[i,0], a0, a1)
    
    if a0<a1:
        a10_list = np.linspace(-0.9,a0-0.01,5)[::-1]
        a01_list = a1-a0+a10_list
    else:
        a01_list = np.linspace(-0.9,a1-0.01,5)[::-1]
        a10_list = a0-a1+a01_list
    ax[i,0].arrow(a0-0.3, a1-0.1, -0.3, -0.3, color='w', width=0.01, head_width=0.05)

    for j, (a10,a01) in enumerate(zip(a10_list,a01_list),1):
        if a0<a1:
            ax[i,j].annotate('$a_{10}$'+'={:2.2}'.format(a10),xy=(0.5,0.2),ha='center',xycoords='axes fraction')
        else:
            ax[i,j].annotate('$a_{01}$'+'={:2.2}'.format(a01),xy=(0.5,0.8),ha='center',xycoords='axes fraction')

        A = np.array([[a0,a01],[a10,a1]])
        flow = partial(ed.lotka_volterra, r=r, a=A)
        traj, info = scipy.integrate.odeint(flow, [B/2,B/2], tspan, full_output=True, mxstep=200 )
        
        # Remove the points after reaching a max pop size. 
        mx = 10
        lower = min(np.min(np.where(traj[:,0]>mx)) if traj[:,0].max()>mx else len(tspan),
                    np.min(np.where(traj[:,1]>mx)) if traj[:,1].max()>mx else len(tspan))
        if mx < len(tspan):
            traj[lower:,0] = np.nan
            traj[lower:,1] = np.nan
            ax[i,j].set(ylim = (0,np.nan_to_num(traj).max()))
        #if info['message'] != "Integration successful.":
            #for k, v in info.items():
            #    print(k,v)
            #print(traj[:,0])
        ax[i,j].plot(tspan, traj[:,0], color='C3')
        ax[i,j].plot(tspan, traj[:,1], color='C0')
        ax[i,j].hlines([1/a0, 1/a1], 0,tspan.max(), colors=['C3','C0'], linestyle='--')
        ax[i,0].scatter(a10,a01,color='white',marker='o', s=50)
        print((a0*a1)/(a10*a01), a10,a01,)
        ax[i,j].set(xlabel='Time', xticks=[], 
                    #title=traj[-1,0]/traj[-1,:].sum(),
                    yticks=[1/a0,B,1/a1],
                    yticklabels=['$a_{00}^{-1}$','B','$a_{11}^{-1}$'])
plt.tight_layout()
plt.savefig('reviews/asymmetry_negative.png', bbox_inches='tight')
plt.savefig('reviews/asymmetry_negative.pdf', bbox_inches='tight')

# %% [code]
fig, ax = plt.subplots(1, 4, figsize=(30,5))

a0,a1 = (0.5,0.5)
a01 = -0.51
a10 = a01
B = 0.00001
A = np.array([[a0,a01],[a10,a1]])
tspan = np.linspace(0,15,10000)
import warnings 
warnings.filterwarnings('error')
 
for j, r in enumerate([[1,1],[1,1.1],[1,1.5],[1,2]]):
    flow = partial(ed.lotka_volterra, r=r, a=A)
    for u in np.linspace(0,1):
        ini = [B*u, (1-u)*B]
        traj = np.zeros((len(tspan),2))
        traj[0,:] = ini
        for u,_ in enumerate(tspan[:-1]):
            try: 
                traj[u+1,:] = traj[u,:] + tspan[1]*flow(traj[u,:])
                if traj[u+1,:].sum()>1000:
                    raise Warning
            except Warning:
                break
        ax[j].plot(tspan[:u], traj[:u,0]/(traj[:u,0]+traj[:u,1]), color='k')
    ax[j].set(ylim=(0,1), xlim=(0,tspan.max()))
    ax[j].set(xlabel='Time within a single generation', title='Growth rates: {}'.format(r))
ax[0].set(ylabel='Collective Colour')
plt.savefig('mutualist_dynamics_within_generation_with_detA_negative.pdf')
```
