## Supplementary material for "Eco-evolutionary dynamics of nested Darwinian populations and the emergence of community-level heredity": Python code, figures and simulations: small_trees.ipynb

```
# %% [markdown]
# Tree View 
This notebook uses the pycairo library to offer a tree-like representation of the data.

# %% [code]
import full_trajectory
import multiprocessing

from functools import partial
import numpy as np
import matplotlib.pyplot as plt
from mpl_toolkits.axes_grid1 import make_axes_locatable
from matplotlib.colors import ListedColormap
from collections import defaultdict
import matplotlib.gridspec as gridspec
import numpy as np

import estaudel.escaffolding as escaffolding
import estaudel.stochastic as stochastic
import estaudel.heredity.process as ehp
import estaudel.heredity.deterministic as ehd
import estaudel.heredity.stochastic as ehs
import estaudel.heredity.treeview as tv

import sys
import PIL.Image as pImage
from PIL.ImageDraw import Draw
import PIL.ImageFont


import os 
from IPython.display import display, Image

from config import SUPFIGPATH, PANEL_DICT

%load_ext autoreload
%autoreload 2

pool = multiprocessing.Pool(6)

# %% [markdown]
# Supplementary figure showcasing the different reproduction regime.

# %% [code]
# Small simulations
np.random.seed(120292)
out = {}
styles = ('serial','neutral','rank','soft')
p = {k:v[0] for k,v in full_trajectory.PARAMETERS.items()}
for style,label in zip(styles,'ABCD'):
    print(style)
    p.update({'N':2,'D':40, 'selection':style, 'percentile':40,
              'collectiveSelectionStrength':0.2,
              'mutation_rate':0,
              'carrying_capacity':200,
           'initial_type0': (0,5,0.6,0.5),
          'initial_type1':(1,5,0.8,0.5)}
        )
    out[style] = full_trajectory.main(p, pool=pool)
    out[style].parents = out[style].parents[:-1]
    t = tv.extract_tree( out[style])
    idx = {n:int(n.split('D')[1]) for n in t['position'].keys()}
    fig = tv.draw_tree_rings(branches=t['branches'],
                             xinfo=t['generation'], 
                             yinfo= t['position'], 
                             zinfo=t['colour'], 
                             oinfo=None,
                             #width=1000, height=6000, size=60)
                             width=333, height=2000, size=20)    
    fname = os.path.join(SUPFIGPATH,'1gen{}.png'.format(style))
    fig.write_to_png(fname)

# %% [code]
def concatenate_images(paths):
    images = [pImage.open(path) for path in paths]
    widths, heights = zip(*(i.size for i in images))
    y_offset = 50
    font_size = y_offset*2
    total_width = sum(widths) + 60
    max_height = max(heights) + y_offset


    new_im = pImage.new('RGBA', (total_width, max_height), color=(0,0,0,1))
    draw = Draw(new_im) 

    x_offset = 10
    tpos = []
    for n,im in enumerate(images):
        new_im.paste(im, (x_offset,0))
        tpos.append(x_offset+im.size[0]/2-font_size/2)
        x_offset += im.size[0] + 20
        if n != len(images)-1:
            draw.line((x_offset, 0, x_offset, max_height), fill='black', width=1)
        x_offset += 20
    new_im = new_im.rotate(-90, expand=True)
    draw = Draw(new_im) 
    ft = PIL.ImageFont.truetype("FreeSans.ttf", font_size)
    for pos,k in zip(tpos,'ABCDEFGHIJKLMNOPQRSTUVWXYZ'):
        draw.text((0,pos),k, fill='black', font=ft)
    return new_im

# %% [code]
styles = ('neutral','rank')
files = [os.path.join(SUPFIGPATH,'1gen{}.png'.format(style)) for style in styles]
print(files)
fig = concatenate_images(files)
fname =  os.path.join(SUPFIGPATH,'1gen_compare.png'.format(style))
fig.save(fname)
Image(fname)

# %% [code]
styles = ('serial','neutral','rank','soft')
fig = concatenate_images( [os.path.join(SUPFIGPATH,'1gen{}.png'.format(style)) for style in styles])
fname =  os.path.join(SUPFIGPATH,'1gen_compare_full.png'.format(style))
fig.save(fname)
Image(fname)

# %% [markdown]
# Small trees

# %% [code]
np.random.seed(120293)
p.update({'N':30,'D':20, 'selection':'neutral', 'percentile':10, 
          'initial_type0': (0,5,0.6,.5),
      'initial_type1':(1,5,0.8,.5)}
    )
out = full_trajectory.main(p, pool=pool)

t = tv.extract_tree(out)
fig = tv.draw_tree_rings(branches=t['branches'],
                         xinfo=t['generation'], 
                         yinfo=t['position'], 
                         zinfo=t['colour'], 
                         oinfo='coal', 
                         width=1000, height=400, size=6, line_width=1
                         #width=3000, height=1200, size=18, line_width=3
                        )
fname_no = os.path.join(SUPFIGPATH,'tree_noselect.png')
fig.write_to_png(fname_no)
Image(filename=fname_no) 

# %% [code]
np.random.seed(120293)
p.update({'N':30,'D':20, 'selection':'rank', 'percentile':10, 
          'initial_type0': (0,5,0.6,.5),
      'initial_type1':(1,5,0.8,.5)}
    )
out = full_trajectory.main(p, pool=pool)

t = tv.extract_tree(out)
fig = tv.draw_tree_rings(branches=t['branches'],
                         xinfo=t['generation'], 
                         yinfo=t['position'], 
                         zinfo=t['colour'], 
                         width=1000, height=400, size=6, line_width=1,
                         #width=3000, height=1200, size=18,
                         oinfo='coal',)
fname_sel = os.path.join(SUPFIGPATH,'tree_select.png')
fig.write_to_png(fname_sel)
Image(filename=fname_sel) 

# %% [code]
# Merge the two trees and add a time axis. 
SVG = """<?xml version="1.0" encoding="UTF-8" standalone="no"?>
<!-- Created with Inkscape (http://www.inkscape.org/) -->

<svg
   xmlns:dc="http://purl.org/dc/elements/1.1/"
   xmlns:cc="http://creativecommons.org/ns#"
   xmlns:rdf="http://www.w3.org/1999/02/22-rdf-syntax-ns#"
   xmlns:svg="http://www.w3.org/2000/svg"
   xmlns="http://www.w3.org/2000/svg"
   xmlns:xlink="http://www.w3.org/1999/xlink"
   xmlns:sodipodi="http://sodipodi.sourceforge.net/DTD/sodipodi-0.dtd"
   xmlns:inkscape="http://www.inkscape.org/namespaces/inkscape"
   width="841.7149mm"
   height="709.93317mm"
   viewBox="0 0 841.7149 709.93317"
   version="1.1"
   id="svg8"
   inkscape:version="0.92.4 (5da689c313, 2019-01-14)"
   sodipodi:docname="tree_supl.svg">
  <defs
     id="defs2">
    <marker
       inkscape:stockid="Arrow2Mend"
       orient="auto"
       refY="0"
       refX="0"
       id="Arrow2Mend"
       style="overflow:visible"
       inkscape:isstock="true">
      <path
         id="path4792"
         style="fill:#000000;fill-opacity:1;fill-rule:evenodd;stroke:#000000;stroke-width:0.625;stroke-linejoin:round;stroke-opacity:1"
         d="M 8.7185878,4.0337352 -2.2072895,0.01601326 8.7185884,-4.0017078 c -1.7454984,2.3720609 -1.7354408,5.6174519 -6e-7,8.035443 z"
         transform="scale(-0.6)"
         inkscape:connector-curvature="0" />
    </marker>
    <marker
       inkscape:stockid="Arrow1Lend"
       orient="auto"
       refY="0"
       refX="0"
       id="Arrow1Lend"
       style="overflow:visible"
       inkscape:isstock="true">
      <path
         id="path4768"
         d="M 0,0 5,-5 -12.5,0 5,5 Z"
         style="fill:#000000;fill-opacity:1;fill-rule:evenodd;stroke:#000000;stroke-width:1.00000003pt;stroke-opacity:1"
         transform="matrix(-0.8,0,0,-0.8,-10,0)"
         inkscape:connector-curvature="0" />
    </marker>
    <marker
       inkscape:stockid="Arrow2Mend"
       orient="auto"
       refY="0"
       refX="0"
       id="Arrow2Mend-7"
       style="overflow:visible"
       inkscape:isstock="true">
      <path
         inkscape:connector-curvature="0"
         id="path4792-5"
         style="fill:#000000;fill-opacity:1;fill-rule:evenodd;stroke:#000000;stroke-width:0.625;stroke-linejoin:round;stroke-opacity:1"
         d="M 8.7185878,4.0337352 -2.2072895,0.01601326 8.7185884,-4.0017078 c -1.7454984,2.3720609 -1.7354408,5.6174519 -6e-7,8.035443 z"
         transform="scale(-0.6)" />
    </marker>
  </defs>
  <sodipodi:namedview
     id="base"
     pagecolor="#ffffff"
     bordercolor="#666666"
     borderopacity="1.0"
     inkscape:pageopacity="0.0"
     inkscape:pageshadow="2"
     inkscape:zoom="0.24748737"
     inkscape:cx="2269.1797"
     inkscape:cy="1202.554"
     inkscape:document-units="mm"
     inkscape:current-layer="layer1"
     showgrid="false"
     fit-margin-top="0"
     fit-margin-left="0"
     fit-margin-right="0"
     fit-margin-bottom="0"
     inkscape:window-width="1920"
     inkscape:window-height="1057"
     inkscape:window-x="0"
     inkscape:window-y="23"
     inkscape:window-maximized="1"
     showguides="false" />
  <metadata
     id="metadata5">
    <rdf:RDF>
      <cc:Work
         rdf:about="">
        <dc:format>image/svg+xml</dc:format>
        <dc:type
           rdf:resource="http://purl.org/dc/dcmitype/StillImage" />
        <dc:title></dc:title>
      </cc:Work>
    </rdf:RDF>
  </metadata>
  <g
     inkscape:label="Layer 1"
     inkscape:groupmode="layer"
     id="layer1"
     transform="translate(431.89069,269.20831)">
    <image
       sodipodi:absref="{fname_no}"
       y="-269.20831"
       x="-390.07144"
       id="image4727"
       xlink:href="{fname_no}"
       preserveAspectRatio="none"
       height="317.5"
       width="793.75" />
    <image
       sodipodi:absref="{fname_sel}"
       y="73.238098"
       x="-390.07144"
       id="image4738"
       xlink:href="{fname_sel}"
       preserveAspectRatio="none"
       height="317.5"
       width="793.75" />
    <text
       xml:space="preserve"
       style="font-style:normal;font-variant:normal;font-weight:normal;font-stretch:normal;font-size:52.91666794px;line-height:1.25;font-family:FreeSans;-inkscape-font-specification:FreeSans;font-variant-ligatures:normal;font-variant-caps:normal;font-variant-numeric:normal;font-feature-settings:normal;text-align:start;letter-spacing:0px;word-spacing:0px;writing-mode:lr-tb;text-anchor:start;fill:#000000;fill-opacity:1;stroke:none;stroke-width:0.90319192"
       x="-428.84735"
       y="-91.170189"
       id="text4743"><tspan
         sodipodi:role="line"
         id="tspan4741"
         x="-428.84735"
         y="-91.170189"
         style="font-style:normal;font-variant:normal;font-weight:normal;font-stretch:normal;font-family:FreeSans;-inkscape-font-specification:FreeSans;stroke-width:0.90319192">A</tspan></text>
    <text
       xml:space="preserve"
       style="font-style:normal;font-variant:normal;font-weight:normal;font-stretch:normal;font-size:52.91666794px;line-height:1.25;font-family:FreeSans;-inkscape-font-specification:FreeSans;font-variant-ligatures:normal;font-variant-caps:normal;font-variant-numeric:normal;font-feature-settings:normal;text-align:start;letter-spacing:0px;word-spacing:0px;writing-mode:lr-tb;text-anchor:start;fill:#000000;fill-opacity:1;stroke:none;stroke-width:0.90319192"
       x="-429.85275"
       y="251.27623"
       id="text4743-3"><tspan
         sodipodi:role="line"
         id="tspan4741-6"
         x="-429.85275"
         y="251.27623"
         style="font-style:normal;font-variant:normal;font-weight:normal;font-stretch:normal;font-family:FreeSans;-inkscape-font-specification:FreeSans;stroke-width:0.90319192">B</tspan></text>
    <path
       style="fill:#000000;stroke:#000000;stroke-width:3;stroke-linecap:butt;stroke-linejoin:miter;stroke-miterlimit:4;stroke-dasharray:none;stroke-opacity:1;marker-end:url(#Arrow2Mend)"
       d="M -390.07144,57.816692 H 403.67855"
       id="path4763"
       inkscape:connector-curvature="0" />
    <path
       style="fill:#000000;stroke:#000000;stroke-width:3;stroke-linecap:butt;stroke-linejoin:miter;stroke-miterlimit:4;stroke-dasharray:none;stroke-opacity:1;marker-end:url(#Arrow2Mend-7)"
       d="m -390.07144,400.26313 h 793.75"
       id="path4763-3"
       inkscape:connector-curvature="0" />
    <text
       xml:space="preserve"
       style="font-style:normal;font-variant:normal;font-weight:normal;font-stretch:normal;font-size:28.22222137px;line-height:1.25;font-family:'Latin Modern Sans';-inkscape-font-specification:'Latin Modern Sans, Normal';font-variant-ligatures:normal;font-variant-caps:normal;font-variant-numeric:normal;font-feature-settings:normal;text-align:start;letter-spacing:0px;word-spacing:0px;writing-mode:lr-tb;text-anchor:start;fill:#000000;fill-opacity:1;stroke:none;stroke-width:0.26458332"
       x="-118.67012"
       y="434.91107"
       id="text5648"><tspan
         sodipodi:role="line"
         id="tspan5646"
         x="-118.67012"
         y="434.91107"
         style="stroke-width:0.26458332">Collective generations</tspan></text>
  </g>
</svg>
""".format(fname_no=os.path.abspath(fname_no), fname_sel=os.path.abspath(fname_sel))

svg_path = os.path.abspath(os.path.join(SUPFIGPATH,'trees.svg'))
with open(svg_path,'w') as file:
    file.write(SVG)
pdf_path = os.path.abspath(svg_path.replace('svg','pdf'))

# %% [code]
# Convert the merged svg to a pdf (Requires inkscape)
!inkscape -without-gui --file=$svg_path --export-pdf=$pdf_path --export-area-snap
```
