## Supplementary material for "Eco-evolutionary dynamics of nested Darwinian populations and the emergence of community-level heredity": Python code, figures and simulations: stats.ipynb

```
# %% [markdown]
# Statistics
Average outcome of several simulations, Figures will go in appendix 1. The data can be gererated using `statistics.condor`. 

# %% [code]
import estaudel.escaffolding as escaffolding
import glob
import pandas as pd
import matplotlib.pyplot as plt
from collections import Counter
import estaudel.heredity.process
import numpy as np
from config import PANEL_DICT

# %% [code]
# Load the data from the multiple results. 
out = []
for f in glob.glob('out/*.csv'):
    out.append(pd.read_csv(f))
    if out[-1].goal.iloc[0]==0.9:
        del out[-1]

st = pd.read_csv('statistics.csv')
out.append(st[st.goal==0.9])

out  = pd.concat(out)
out.columns = [x if x[0]!='(' else eval(x) for x in out.columns]
out.initial_type0 = [x if x[0]!='(' else eval(x) for x in out.initial_type0]
out.initial_type1 = [x if x[0]!='(' else eval(x) for x in out.initial_type1]

out['extinct'] = [(mcp==1 or mcp==0)  for mcp in out['mean_cp']]
out['mean_deviation_to_target_colour'] = out.mean_cp - (1-out.goal)
out['mean_deviation_to_target_colour_percent'] = out['mean_deviation_to_target_colour'] * 100

columns = ['mean_cp', ('red', 'growth_rate'),
        ('red', 'interaction_inter'),       ('blue', 'growth_rate'),
       ('blue', 'interaction_inter')]

assert out.N.nunique() == 1

print("Loaded the endpoint of {} simulations".format(out.shape[0]))

# %% [code]
def violin(keys, distrib, ax):
    ax.violinplot(distrib, showmeans=True)
    ax.set(xticks=range(1,len(keys)+1),
           xticklabels=[str(x) for x in keys])
    ax.set_xticklabels([str(x) for x in keys], rot=90)
    
def get_param(v):
    assert len(v.initial_type0.unique()) == 1
    assert len(v.initial_type1.unique()) == 1
    return dict(
    r0 = v.initial_type0.unique()[0][1],
    r1 = v.initial_type1.unique()[0][1],
    a00 = v.initial_type0.unique()[0][2],
    a11 = v.initial_type1.unique()[0][2],
    a01_i = v.initial_type0.unique()[0][3],
    a10_i = v.initial_type1.unique()[0][3])

def plot_violins(cc, data, ax, title=None):
    for i, c in enumerate(cc): 
        keys, distrib, df = zip(*[(k, v[c].values, v) for k, v in data])
        pos = np.arange(len(keys))*2+i*0.5
        parts = ax.violinplot(distrib, positions=pos, showmeans=True)
        try:
            keys = ["$a_{{red}}^{{intra}}$ = {}".format(k[2]) for k in keys]
        except Exception:
            keys = ["{}".format(k) for k in keys]
        for k, x, d, di in zip(keys, pos, df, distrib):
            #ax.text(x, np.max(di)+0.01, '{}/{}'.format(d.shape[0]-d['extinct'].sum(),d.shape[0]),
            #       verticalalignment='bottom', horizontalalignment='center'
            #       )
            if 'interaction' in title:
                param = get_param(d)
                pred = (1-d.goal.unique()[0])*param['a11']-param['a00']
                if pred>0:
                    if i:
                        ax.hlines(abs(pred), x-0.25,x+0.25,  color='b', ls='--')
                    else:
                        ax.hlines(0, x-0.25,x+0.25,  color='r', ls='--')
                else:
                    if not i:
                        ax.hlines(abs(pred), x-0.25,x+0.25,  color='r', ls='--')
                    else:
                        ax.hlines(0, x-0.25,x+0.25,  color='b', ls='--')

        for pc in parts['bodies']:
            pc.set_facecolor(c[0])
        for pc in [parts['cmeans'],parts['cmaxes'],parts['cmins'],parts['cbars']]:
            pc.set_edgecolor(c[0])

    ax.set(title=title)
    ax.set(xticks=pos-0.25, xticklabels=keys)
    #ax.set(yticks=[0,0.25,0.4], yticklabels=[0,0.25,0.4])

# %% [code]
plt.rc('font', size=15)
fig, axes= plt.subplots(1,3, figsize=(20,5))

for k,ax in enumerate(axes):
    ax.annotate('ABCD'[k],**PANEL_DICT)

xt = [-0.6,-0.4,-0.2,0,0.2,0.4,0.6]
axes[0].set(yticks=xt, ylim=(-0.75,0.75), yticklabels=["{}%".format(x) for x in xt])
plot_violins([ ('mean_deviation_to_target_colour_percent')],
               out[out.goal==0.5].groupby(['initial_type0']), ax=axes[0],
            title='Mean deviation to optimal colour')
plot_violins([ ('red', 'interaction_inter'), ('blue', 'interaction_inter')],
               out[out.goal==0.5].groupby(['initial_type0']), ax=axes[1],
            title='Inter-colour competition')
plot_violins([ ('red', 'growth_rate'), ('blue', 'growth_rate')],
             out[out.goal==0.5].groupby(['initial_type0']), ax=axes[2],
            title='Growth rates')
axes[1].set(ylim=(-0.01,0.35))
plt.savefig('supfig/traits_reproducibility.pdf',  bbox_inches='tight')

# %% [code]
idx = out.initial_type0 == (0, 5, 0.7, 0.3)
fig, axes= plt.subplots(1,3, figsize=(20,5))
plot_violins([ ('mean_deviation_to_target_colour_percent')],
               out[idx].groupby(['goal']), ax=axes[0],
            title='Mean deviation to optimal colour')
xt = np.arange(-6,7,2)
axes[0].set(yticks=xt, ylim=(-5,5), yticklabels=["{}%".format(x) for x in xt])

for k,ax in enumerate(axes):
    ax.annotate('ABCD'[k],**PANEL_DICT)

#axes[0].hlines(,0,10, color='k', ls='--')
plot_violins([ ('red', 'interaction_inter'), ('blue', 'interaction_inter')],
               out[idx].groupby(['goal']), ax=axes[1],
            title='Interaction parameters')
plot_violins([ ('red', 'growth_rate'), ('blue', 'growth_rate')],
             out[idx].groupby(['goal']), ax=axes[2],
            title='Growth rates')

for ax in axes:
    ax.set(xlabel='Optimal colour $\hat \phi$')
plt.savefig('supfig/traits_different_objectives.pdf',  bbox_inches='tight')

# %% [code]
import estaudel.heredity.deterministic as ehd

# %% [code]
plt.rc('font', size=15)
fig, axes= plt.subplots(1,2, figsize=(12,5))
for k,ax in enumerate(axes):
    ax.annotate('ABCD'[k],**PANEL_DICT)
axes[0].set(xlim=(0,.4), ylim=(0,.4),
            xlabel='$a_{red}^{inter}$',
            ylabel='$a_{blue}^{inter}$',
            title='Inter-colour competition',
            xticks = [0,.1,.2,.3,.4], 
            yticks = [0,.1,.2,.3,.4])
axes[1].set(xlim=(0,20),ylim=(0,20), 
            xticks = [0,5,10,15,20],
            yticks=[0,5,10,15,20],
            xlabel='$r_{red}$', 
            title='Growth rates',
            ylabel='$r_{blue}$')
axes[0].scatter(0.3, 0.3, color='k', label='Initial conditions')
axes[1].scatter(5, 5, color='k', label='Initial conditions')

for (goal,it0),v in  out[out.goal==0.5].groupby(['goal','initial_type0']):
    x = np.linspace(0,1)
    _,r0, a00, a01 = v['initial_type0'].iloc[0] 
    _,r0, a11, a10 = v['initial_type1'].iloc[0] 
    marker = '.'
    if a00 == a11:
        lab = r'Equal c. capacity, $a_{red}^{intra}=a_{blue}^{intra}$'
        marker='x'
    elif a11 > a00:
        lab = 'Red higher c. capacity, $a_{red}^{intra}<a_{blue}^{intra}$'
        marker='+'
    elif a00 > a11:
        lab = 'Blue higher c. capacity, $a_{red}^{intra}>a_{blue}^{intra}$'
        marker='.'
    axes[0].plot(a00+(1-1/goal)*(a11-x), x, ls=":")
    axes[0].scatter(v[('red', 'interaction_inter')], v[('blue', 'interaction_inter')], marker=marker)
    axes[1].scatter(v[('red', 'growth_rate')], v[('blue', 'growth_rate')], 
                    marker=marker, label=lab)
axes[1].legend(loc='lower right',  prop={'size': 10})
plt.tight_layout()
plt.savefig('supfig/traits_reproducibility_deterministic.pdf',  bbox_inches='tight')

# %% [code]
plt.rc('font', size=15)
fig, axes= plt.subplots(1,2, figsize=(12,5))
for k,ax in enumerate(axes):
    ax.annotate('ABCD'[k],**PANEL_DICT)
axes[0].set(xlim=(0,1), ylim=(0,1),
            xlabel='$a_{red}^{inter}$',
            ylabel='$a_{blue}^{inter}$',
            title='Inter-colour competition',
)
axes[1].set(xlim=(0,20),ylim=(0,20), 
            xticks = [0,5,10,15,20],
            yticks = [0,5,10,15,20],
            xlabel = '$r_{red}$', 
            title = 'Growth rates',
            ylabel='$r_{blue}$')
axes[0].scatter(0.3, 0.3, color='k', label='Initial conditions')
axes[1].scatter(5, 5, color='k', label='Initial conditions')

for k,v in  out[out['initial_type0']==(0, 5, 0.7, 0.3)].groupby('goal'):
    x = np.linspace(0,1)
    _,r0, a00, a01 = v['initial_type0'].iloc[0] 
    goal = k 
    _,r0, a11, a10 = v['initial_type1'].iloc[0] 
    l = axes[0].plot(a00+(1-1/goal)*(a11-x), x, ls=":")
    u = 0.02+goal*0.7
    axes[0].text(u,(a11-(u-a00)/(1-1/goal)), '${}$'.format(goal), color=l[0].get_color())
    axes[0].scatter(v[('red', 'interaction_inter')], v[('blue', 'interaction_inter')], marker='.')
    axes[1].scatter(v[('red', 'growth_rate')], v[('blue', 'growth_rate')], 
                    label=lab,
                    marker='.')
plt.tight_layout()
plt.savefig('supfig/traits_different_objective_deterministic.pdf', bbox_inches='tight')

# %% [code]
```
