## Supplementary material for "Eco-evolutionary dynamics of nested Darwinian populations and the emergence of community-level heredity": Python code, figures and simulations: Supplementary_figures.ipynb

```
# %% [markdown]
# Supplementary figures

# %% [code]
from collections import OrderedDict, Counter
from multiprocessing import Pool
from functools import partial
import pickle

import numpy as np

import scipy.optimize
import scipy.integrate
import scipy.misc

# Matplotlib related imports 
import matplotlib.pyplot as plt
from matplotlib import rc
from matplotlib.colors import LinearSegmentedColormap
from matplotlib.colors import ListedColormap
from mpl_toolkits.axes_grid1.axes_divider import make_axes_locatable
from mpl_toolkits.axes_grid1.colorbar import colorbar
from mpl_toolkits.axes_grid1.inset_locator import inset_axes, zoomed_inset_axes
from mpl_toolkits.axes_grid1.anchored_artists import AnchoredSizeBar
%matplotlib inline

import estaudel.escaffolding as escaffolding
import estaudel.heredity.process
import estaudel.heredity.deterministic as ehd
from estaudel.heredity.stochastic import COLOR_NAMES
from config import BLUE_RED, PANEL_DICT

# %% [markdown]
# Parameter sets

# %% [code]
Akeys = ['0excludes1', '1excludes0', 'bistable', 'coex']
rkeys = ['0fast', '1fast']

Aset = {
 'coex': np.array([[0.7, 0.3],
                 [0.5, 0.6]]),
 '1excludes0':np.array([[0.6, 0.7],
                      [0.4, 0.3]]),
 '0excludes1':np.array([[0.3, 0.6],
                       [0.5, 0.8]]),
 'bistable':np.array([[0.3, .7],
                      [0.8, 0.2]]),
}

r_base = 30
rset = {'0fast':np.array([1.2,1])*r_base,
        '1fast':np.array([1,1.2])*r_base}

B = 0.001

rlabel = {'1fast':COLOR_NAMES[1].capitalize()+' fast grower','0fast':COLOR_NAMES[0].capitalize()+' fast grower'}
alabel = {'bistable':'Bistable', 'coex':'Stable Coexistence', 
          '0excludes1':COLOR_NAMES[1].capitalize()+' extinction',
          '1excludes0':COLOR_NAMES[0].capitalize()+' extinction'}

alabel = {'bistable':'Bistable', 'coex':'Stable Coexistence', 
          '0excludes1':COLOR_NAMES[1].capitalize()+' extinction',
          '1excludes0':COLOR_NAMES[0].capitalize()+' extinction'}

alabel={ 'coex': "Asymptotic Coexistence",
 '1excludes0': "Competitive exclusion of the fast grower",
 '0excludes1': "Competitive exclusion of the slow grower",
 'bistable': "Asymptotic Bistability",}

# %% [markdown]
# Stream plots in N0,N1 coordinates.

This figure shows the vector field of the Density-dependent ecology in the (N0,N1) coordinate space for four sets of parameters representative of the four qualitatively different behaviors.

# %% [code]
def plot_vector_field(r, A, color='grey', text=True, ax=None, mx=None, B=None):
    """Plot the LV-vector field with null isoclines and equilibrium."""
    if ax is None:
        ax = plt.gca()
        
    # Flow 
    if mx is None:
        mx = np.max(1/A)
    dx = mx/30
    x = np.linspace(0,mx+dx,20)
    X,Y = np.meshgrid(x,x)
    
    flow = np.vectorize(lambda x,y: ehd.lotka_volterra([x,y],r=r,a=A),
                        signature='(),()->(2)')
    U = flow(X,Y)
    ax.streamplot(X, Y, U[:,:,0], U[:,:,1], density=1, color=color,linewidth=0.8)
    
    # Trivial null-isoclines
    ax.plot([0,mx],[0,0], color='b', ls='-', alpha=0.3, lw=5)
    ax.plot([0,0],[mx,0], color='r', ls='-', alpha=0.3, lw=5)

    # Non trivial null isoclines
    ax.plot([0, 1/A[1,0]], [1/A[1,1], 0], color='b', ls='-', alpha=0.3, lw=5)
    ax.plot([0, 1/A[0,0]], [1/A[0,1], 0], color='r', ls='-', alpha=0.3, lw=5)
    
    # Initial conditions
    if B is not None:
        ax.plot([0,B],[B,0], color='C4', ls='-', alpha=0.3, lw=5)
    
    # Equilibria 
    for e,s in ehd.list_equilibria(A):
        ax.scatter(*e, color='k' if s else 'w', edgecolor='w' if s else 'k')
        
    if False:
        ax.text(1/A[0,0], 0, r'$a_{00}^{-1}$', color='r',
            horizontalalignment='center', verticalalignment='top')
        ax.text(0, 1/A[1,1], r'$a_{11}^{-1}$', color='b',
            horizontalalignment='right', verticalalignment='center')
    else:
        ax.set(xticks = [0, 1/A[0,0]],
               xticklabels=[0,r'$a_{00}^{-1}$'])
        ax.set(yticks = [0, 1/A[1,1]],
               yticklabels=[0,r'$a_{11}^{-1}$'])
    
    ax.set(xlabel='$N_0$',ylabel='$N_1$',
           #xticks=[],yticks=[],
           xlim=(-dx,mx+dx), ylim=(-dx,mx+dx))

# %% [code]
f, axes = plt.subplots(2,2,figsize=(8,8))
for k,ax in zip(Akeys, axes.flat):
    plot_vector_field(rset['0fast'],Aset[k], ax=ax)
    ax.set(title=alabel[k])
plt.tight_layout()
plt.savefig('supfig/paramspace_streamplot.pdf', bbox_inches='tight')

# %% [markdown]
# Coordinates

This figure show the phase diagram in both (N0,N1) coordinate and (N,µ) coordinates.

# %% [code]
def isocline_nontriviale_N(f,r,a):
    """Non trivial null-isocline for N in Nf coordinates."""
    return (r[0]*f - r[1]*f + r[1])/(a[0,0]*r[0]*f**2 - a[0,1]*r[0]*f**2 + a[0,1]*r[0]*f - a[1,0]*r[1]*f**2 + a[1,0]*r[1]*f + a[1,1]*r[1]*f**2 - 2*a[1,1]*r[1]*f + a[1,1]*r[1])
def isocline_nontrivale_f(N,r,a):
    """Non trivial null-isocline for f in Nf coordinates. """
    return (-a[0,1]*r[0]*N + a[1,1]*r[1]*N + r[0] - r[1])/((a[0,0]*r[0] - a[0,1]*r[0] - a[1,0]*r[1] + a[1,1]*r[1])*N)

def plot_nf_vectorfield(r, A, color='grey', ax=None, xmax=2.5, text=True, B=None):
    """Plot the LV-vector field in NF coordinates."""
    if ax is None:
        ax = plt.gca()
    margin_y = 0.01
    y = np.linspace(-margin_y,1+margin_y,20)
    x = np.linspace(0,xmax,20)

    # Isoclines
    ntN = isocline_nontriviale_N(y, r, A)
    ax.plot(ntN,y,color='orange', ls='-',alpha=0.3,lw=5)

    ntf = isocline_nontrivale_f(x, r,A)
    ax.plot(x,ntf,color='green', ls='-',alpha=0.3,lw=5)

    ax.plot([0,x.max()],[0,0], color='g', ls='-',alpha=0.3,lw=5)
    ax.plot([0,x.max()],[1,1], color='g', ls='-',alpha=0.3,lw=5)
    ax.plot([0,0],[0,1], color='orange', ls='-',alpha=0.3,lw=5)
    
    # Initial conditions
    if B is not None:
        ax.plot([B,B],[0,1], color='C4', ls='-', alpha=0.3, lw=5)
        
    if False:
        ax.text(1/A[0,0], 1, r'$a_{00}^{-1}$', color='k',
                horizontalalignment='center', verticalalignment='bottom')
        ax.text(1/A[1,1], 0, r'$a_{11}^{-1}$', color='k',
                horizontalalignment='center', verticalalignment='top')

    # Equilibria
    for e,s in ehd.list_equilibria(A):
        N = e[0]+e[1]
        if N:
            f = e[0]/N
            ax.scatter(N,f, color='k' if s else 'w', edgecolor='w' if s else 'k')

    # Flow 
    X,Y = np.meshgrid(x,y)
    flow = np.vectorize(lambda x,y: ehd.lotka_volterra_Nf([x,y],r=r,a=A),
                        signature='(),()->(2)')
    U = flow(X,Y)
    ax.streamplot(X, Y, U[:,:,0], U[:,:,1], density=1.2, color=color,linewidth=0.8)
    ax.set(xlabel='Population size, N',ylabel='Collective colour, x',
           ylim=(-margin_y,1+margin_y),
           xlim=(0,x.max()),yticks=[0,0.5,1])
    
    ax.set(xticks=[0,1/A[1,1]],
           xticklabels=[0,r'$a_{11}^{-1}$'])
    ax2 = ax.twiny()
    ax2.set(xticks=[1/A[0,0]],
            xlim=(0,x.max()),
            xticklabels=[r'$a_{00}^{-1}$'])

# %% [code]
plt.rc('font', size=15)
plt.figure(figsize=(13,5))

ax1 = plt.subplot2grid((1,3), (0,0), rowspan=1, colspan=1)
plot_vector_field(rset['0fast'], Aset['coex'], 'grey', ax=ax1)


ax2 = plt.subplot2grid((1,3), (0,1), rowspan=1, colspan=2)
plot_nf_vectorfield(rset['0fast'], Aset['coex'], 'grey', ax=ax2)
plt.tight_layout()
plt.savefig('supfig/nf_coordinate_streamplot.pdf',bbox_inches='tight')

# %% [markdown]
# Bifurcation diagrams

This figure show the bifurcation diagram for $µ_{eq}$ on parameter $T$

# %% [code]
def plot_bifurcation_diagram(title, pstar, tstar, stability_of_01, continuation, stability_of_continuation,
                             color='C0', ax=None, plot_cont_info=False, **_):
    """Display bifurcation diagram"""
    
    if ax is None:
        ax = plt.gca()
        
    ax.grid(alpha=.1)

    # Draw the 0 and 1 branches
    for k,v in stability_of_01.items():
        for n,seg in enumerate(v):
            ax.hlines(k,seg[1][0],seg[1][1],
                      linestyles='-' if seg[0] else ':',
                      color=color,)
                      #label=('Stable' if seg[0] else 'Instable') if n==0 else None)
    ax.hlines(0,0,0, color=color, linestyles='-', label='Stable')
    ax.hlines(0,0,0, color=color, linestyles=':', label='Instable')

    # Draw the continuation branch
    ax.plot(continuation[0], continuation[1], color='C0',
           ls='-' if stability_of_continuation else ':')
    
    if plot_cont_info and cont_info is not None:
        ax.scatter(cont_info['t0'],cont_info['x0'], color='grey')
        ax.vlines(cont_info['tmin'],0,1, color='grey', alpha=0.1, lw=3)
        ax.vlines(cont_info['tmax'],0,1, color='grey', alpha=0.1, lw=3)
   
    ax.set(title=title,
           xlabel='Duration of the growth phase, T', 
           ylabel=r'Adult collective colour, $\phi$')

# %% [code]
# Compute all the bifurcation diagram
databif = {}
for i,akey in enumerate(Akeys):
    for j,rkey in enumerate(rkeys):
        databif[(i,j)] = {'akey':akey, 'rkey':rkey}
        databif[(i,j)].update(ehd.get_bifurcation_diagram(rset[rkey],Aset[akey],B,tmax=1))

# %% [code]
def h(t, side, r, A, B):
    x,y = (0,1) if side==0 else (1,0)
    beta = A[x,x]*B
    alpha = r[x]*r[y]*(A[x,x]-A[y,x])/(r[x]*A[x,x]-r[y]*A[y,x])
    return beta - (1+beta)*np.exp(-alpha*t)+np.exp(-r[x]*t)

# %% [code]
# Compute all the bifurcation diagram
databif = []
rkey = '0fast'
for i,akey in enumerate(Akeys):
    databif.append({'akey':akey, 'rkey':rkey})
    databif[-1].update(ehd.get_bifurcation_diagram(rset[rkey],Aset[akey],B,tmax=1))

# %% [code]
fig,axes = plt.subplots(2,2,figsize=(15,7.5))

for k,d in enumerate(databif):
    axes.flat[k].annotate('ABCD'[k],**PANEL_DICT)
    d['title']=alabel[d['akey']]
    plot_bifurcation_diagram(ax=axes.flat[k], **d)
    axes.flat[k].legend(loc=6)
    
    val, _, conv, _ = scipy.optimize.fsolve(partial(h, side=1, r=rset[d['rkey']],A=Aset[d['akey']],B=B), 
                                        0.1, full_output=True)
    val2, _, conv, _ = scipy.optimize.fsolve(partial(h, side=0, r=rset[d['rkey']],A=Aset[d['akey']],B=B), 
                                        0.1, full_output=True)
    #val2 = val2 if val2<1 else np.nan
    #val = val if val<1 else np.nan
    
    #print(val, val2)
    axes.flat[k].set(xticks = [val,val2], xticklabels=['$T^*_B$', '$T^*_R$'],xlim=(0,1))
plt.tight_layout()
plt.savefig('supfig/bifurcation_diagrams.pdf',bbox_to_inches='tight')

# %% [markdown]
# p*

# %% [markdown]
This figure show the equilibrium of the dynamical system as a function of the inter-color interaction parameters.

# %% [code]
xx = np.linspace(0,1,200)
Xa,Ya = np.meshgrid(xx,xx)
a0 = 0.7
a1 = 0.8
f = np.vectorize(lambda x,y: estaudel.heredity.deterministic.pstar(np.array([[a0,y],
                                                                             [x,a1]])))
Za = f(Xa,Ya)

ticks = [0,0.25,0.5,0.75,1]
levels = np.linspace(0,1,20)

fig, ax = plt.subplots(1,1, figsize=(8,7))
div = make_axes_locatable(ax)
cax = div.append_axes("right", size="4%", pad=0.11)

# Background 
im = ax.contourf(Xa, Ya, Za, cmap=BLUE_RED, levels=levels)
ax.contourf(Xa, Ya, np.int_(np.isnan(Za)), colors='white', levels=[0.1,1],  hatches='/') 
ax.contour(Xa,Ya,Za,colors='w',linestyles='--',levels=[0.5])

#ax.contourf(Xa, Ya, np.int_(np.isnan(Za)), colors='white', levels=[0.1,1],  hatches='/') 

ax.fill_between(x=[a0,1],y1=[a1,a1], y2=[1,1], color='#bab0ac')

# Colorbar
cb = plt.colorbar(im, cax=cax, ticks=ticks)
ax.hlines(a1,0,1,color='w',linestyles='-')
ax.vlines(a0,0,1,color='w',linestyles='-')
cax.plot([0,1],[0.5,0.5],color='w',ls=':')
cax.plot([0,1],[1,1],color='w',ls='-')
cax.plot([0,1],[0,0],color='w',ls='-')

# Set axis
ax.set(
        xlabel=r'$a_{10}$', 
        ylabel=r'$a_{01}$',
        title=r'Stable fixed point of $G_\theta$ when $T \to \infty$',
        xlim=(Xa[0,0],Xa[-1,-1]),
        ylim=(Ya[0,0],Ya[-1,-1]),
        xticks=[0,a0,1],
        xticklabels=[0,'$a_{00}$',1,],
        yticklabels=[0, '$a_{11}-a_{00}$','$a_{11}$',1,],
        yticks=[0,a1-a0,a1,1]
);
plt.tight_layout()
plt.savefig('supfig/position_of_pstar.pdf')

# %% [code]
def minipstar(ax, a0,a1):
    x = np.linspace(0,a0, 10)
    ax.plot(x, a1-a0+x, color='white')
    ax.fill_between(x=[a0,1],y1=[a1,a1], y2=[1,1], color='#bab0ac')
    ax.fill_between(x=[a0,1],y1=[0,0], y2=[a1,a1], color=BLUE_RED(1.0))
    ax.fill_between(x=[0,a0],y1=[a1,a1], y2=[1,1], color=BLUE_RED(0.0))
    if a0<a1:
        tik = dict(xticks=[0,a0,1],
            xticklabels=[0,'$a_{00}$',1,],
            yticklabels=[0, '$a_{11}-a_{00}$','$a_{11}$',1,],
            yticks=[0,a1-a0,a1,1])
    else:
        tik = dict(xticks=[0,a0-a1,a0,1],
        xticklabels=[0,'$a_{00}-a_{11}$','$a_{00}$',1,],
        yticklabels=[0, '$a_{11}$',1,],
        yticks=[0,a1,1])
    ax.set(xlabel='$a_{10}$', ylabel='$a_{01}$', xlim=(0,1), ylim=(0,1), **tik)
        
    Xa,Ya = np.meshgrid(np.linspace(0,a0-0.001,10),np.linspace(0,a1-0.001,10))
    f = np.vectorize(lambda x,y: estaudel.heredity.deterministic.pstar(np.array([[a0,y],
                                                                             [x,a1]])))
    Za = f(Xa,Ya)
    levels = np.linspace(0,1,20)
    im = ax.contourf(Xa, Ya, Za, cmap=BLUE_RED, levels=levels)

# %% [code]
fig, ax = plt.subplots(2, 6, figsize=(20,6.66))
a = [(0.7, 0.8), 
     (0.8,0.5)]
r = [60,40]
tspan = np.linspace(0,1,1000)
for i, (a0,a1) in enumerate(a):
    minipstar(ax[i,0], a0, a1)
    
    if a0<a1:
        a10_list = np.linspace(0,a0-0.01,5)[::-1]
        a01_list = a1-a0+a10_list
    else:
        a01_list = np.linspace(0,a1-0.01,5)[::-1]
        a10_list = a0-a1+a01_list
    ax[i,0].arrow(a0-0.3, a1-0.1, -0.3, -0.3, color='w', width=0.01, head_width=0.05)


    for j, (a10,a01) in enumerate(zip(a10_list,a01_list),1):
        if a0<a1:
            ax[i,j].annotate('$a_{10}$'+'={:2.2}'.format(a10),xy=(0.5,0.2),ha='center',xycoords='axes fraction')
        else:
            ax[i,j].annotate('$a_{01}$'+'={:2.2}'.format(a01),xy=(0.5,0.8),ha='center',xycoords='axes fraction')

        A = np.array([[a0,a01],[a10,a1]])
        flow = partial(ehd.lotka_volterra, r=r, a=A)
        traj = scipy.integrate.odeint( flow,[B/2,B/2], tspan )
        ax[i,j].plot(tspan, traj[:,0], color='C3')
        ax[i,j].plot(tspan, traj[:,1], color='C0')
        ax[i,j].hlines([1/a0, 1/a1], 0,tspan.max(), colors=['C3','C0'], linestyle='--')
        ax[i,0].scatter(a10,a01,color='white',marker='o', s=50)
        
        ax[i,j].set(xlabel='Time', xticks=[], yticks=[1/a0,B,1/a1], yticklabels=['$a_{00}^{-1}$','B','$a_{11}^{-1}$'])
plt.tight_layout()
plt.savefig('supfig/asymmetry.png', bbox_inches='tight')
plt.savefig('supfig/asymmetry.pdf', bbox_inches='tight')

# %% [code]
fig, ax = plt.subplots(2, 6, figsize=(18,6))
plist = np.linspace(0,1,100)
for i, (a0,a1) in enumerate(a):
    minipstar(ax[i,0], a0, a1)
    if a0<a1:
        a10_list = np.linspace(0,a0-0.01,5)
        a01_list = a1-a0+a10_list
    else:
        a01_list = np.linspace(0,a1-0.01,5)
        a10_list = a0-a1+a01_list
    
    for j, (a10,a01) in enumerate(zip(a10_list,a01_list),1):
        A = np.array([[a0,a01],[a10,a1]])
        g = ehd.get_gfunc(r=r,A=A, B=B, T=1)
        ax[i,j].plot([0,1],[0,1], color='k')
        ax[i,j].plot(plist, [g(p) for p in plist])
        #ax[i,j].set(xlim=(0,1),ylim=(0,1))
        ax[i,0].scatter(a10,a01,color='white',marker='+')
plt.tight_layout()
```
