## Supplementary material for "Eco-evolutionary dynamics of nested Darwinian populations and the emergence of community-level heredity": Python code, figures and simulations: README.md

# estaudel - Nested Darwinian population simulation

## Cite this work

> Eco-evolutionary dynamics of nested Darwinian populations and the emergence of  community-level heredity
> Guilhem Doulcier, Amaury Lambert, Silvia De Monte, Paul B. Rainey
> bioRxiv 827592; doi: https://doi.org/10.1101/827592

## Content:

- `/estaudel/` : is the python library
- `/estaudel/stochastic` : Implement a stochastic birht-death-process with mutation for particles.
- `/estaudel/escafolding` : Implement collective level events, like collective reproduction and selection.
- `/estaudel/heredity`: Implement the red/blue model used in the heredity project.
- `/evolution_of_heredity/` : contains code for the figure in the
  "Eco-evolutionary dynamics of nested Darwinian populations and the
emergence of community-level heredity" article.
- `/evolution_of_heredity/full_trajectory.py`: shows how to use the bits
  and pieces from the library and do a full simulation.

## Getting started

Here is a quick way of setup a development environment, independant from your python install.

```
# 1. Create a folder to store everything.
mkdir nested_evolution
cd nested_evolution

# 2. Get the source
git clone git@gitlab.com:ecoevomath/estaudel.git

# 3. Optional: Create a virtual environment
# Use this if you do not want to mess with the python package installed on your computer.
# You do not need root access, but you will need the virtualenv package
# If you want to install on your main python environment, you can safely jump to step 4.
virtualenv env # `sudo pip3 install virtualenv` will install the package if it is missing.
source env/bin/activate  # Switch to the virtual environment

# 4. Install the python package in "developer mode".
pip3 install -e estaudel/

# 5. Try it out.
python3 estaudel/evolution_of_heredity/full_trajectory.py --N 10 --D 10
```

The figures are produced by code found in the Jupyter notebooks within
the `evolution_of_heredity` folder.

## License

Copyright (C) Guilhem Doulcier 2016-2020

This program is free software: you can redistribute it and/or modify
it under the terms of the GNU General Public License as published by
the Free Software Foundation, either version 3 of the License, or
(at your option) any later version.

This program is distributed in the hope that it will be useful, but
WITHOUT ANY WARRANTY; without even the implied warranty of
MERCHANTABILITY or FITNESS FOR A PARTICULAR PURPOSE.  See the GNU
General Public License for more details.

You should have received a copy of the GNU General Public License
along with this program.  If not, see <http://www.gnu.org/licenses/>.
