## Supplementary material for "Eco-evolutionary dynamics of nested Darwinian populations and the emergence of community-level heredity": Python code, figures and simulations: setup.py

```
from setuptools import setup

setup(name='estaudel',
      version='0.1',
      description='Nested Darwinian population simulations',
      url='https://gitlab.com/ecoevomath/estaudel',
      author='Guilhem Doulcier',
      author_email='',
      license='GPLv3+',
      python_requires='>=3',
      packages=['estaudel'],
      install_requires=[
          'numpy',
          'scipy',
          'pandas',
          'matplotlib',
          'pycairo',
          'Pillow'],
      zip_safe=False)
```
